# A pro-inflammatory stem cell niche drives myelofibrosis through a targetable galectin 1 axis

**DOI:** 10.1101/2023.08.05.550630

**Authors:** Rong Li, Michela Colombo, Guanlin Wang, Antonio Rodriguez-Romera, Jennifer O’Sullivan, Sally-Ann Clark, Juan M. Pérez Sáez, Yiran Meng, Abdullah O. Khan, Sean Wen, Pengwei Dong, Wenjiang Zhou, Nikolaos Sousos, Lauren Murphy, Matthew Clarke, Natalie J. Jooss, Aude-Anais Olijnik, Zoë C. Wong, Christina Simoglou Karali, Korsuk Sirinukunwattana, Hosuk Ryou, Ruggiero Norfo, Qian Cheng, Charlotte K. Brierley, Joana Carrelha, Zemin Ren, Supat Thongjuea, Vijay A Rathinam, Anandi Krishnan, Daniel Royston, Gabriel A. Rabinovich, Adam J Mead, Bethan Psaila

## Abstract

Myeloproliferative neoplasms are stem cell-driven cancers associated with a large burden of morbidity and mortality. The majority of patients present with early-stage disease, but a substantial proportion progress to myelofibrosis and/or secondary leukemia, advanced cancers with a poor prognosis and high symptom burden. Currently, it remains difficult to predict progression, and we lack therapies that reliably prevent or reverse fibrosis development. A major bottleneck to the discovery of disease-modifying therapies has been an incomplete understanding of the interplay between perturbed cellular and molecular states. Several cell types have individually been implicated, but a comprehensive analysis of myelofibrotic bone marrow is lacking. We therefore mapped the crosstalk between bone marrow cell types in myelofibrotic bone marrow. We found that inflammation and fibrosis are orchestrated by a ‘quartet’ of immune and stromal cell lineages – with basophils and mast cells creating a TNF signaling hub, communicating with megakaryocytes, mesenchymal stromal cells and pro-inflammatory fibroblasts. We identified the β-galactoside binding protein galectin 1 as a striking biomarker of progression to myelofibrosis and poor survival in multiple patient cohorts, and as a promising therapeutic target, with reduced myeloproliferation and fibrosis *in vitro* and *in vivo* and improved survival following galectin 1 inhibition. In human bone marrow organoids, TNF increased galectin 1 expression, suggesting a feedback loop wherein the pro-inflammatory MPN clone creates a self-reinforcing niche, fueling progression to advanced disease. This study provides a valuable resource for studying hematopoietic cell-niche interactions, with broad relevance for cancer-associated inflammation and disorders of tissue fibrosis.

## INTRODUCTION

In most cancers, one or more genetic perturbations are initiating events that confer a survival advantage to the cell-of-origin and its progeny, but the stromal-immune context in which the emergent clone operates determines its ultimate impact. Myeloproliferative neoplasms (MPNs) are initiated by somatic mutations in hematopoietic stem cells (HSCs) that cause clonal expansion and an over-production of blood cells and their progenitors (*1*). The underlying genetic lesions are well described, with mutations affecting either the gene encoding the Janus kinase signal transducer JAK2 (JAK2V617F), the chaperone protein calreticulin (CALR) or the thrombopoietin receptor (MPL) occurring in almost all patients (*2*). Interactions between the MPN clone and its microenvironment influence the rate and likelihood of progression to advanced disease (*3–5*). While most patients present with slow-growing malignancies that only modestly impact life expectancy, some patients develop a severe form of MPN called myelofibrosis. In these patients, fibrotic bone marrow remodeling and pronounced systemic inflammation cause bone marrow failure, extramedullary hematopoiesis, splenomegaly, severe symptoms and a median survival of around 5 years (*6*).

Myelofibrosis results when cytokines produced by the MPN clone stimulate bone marrow stromal cells to deposit an excess of collagens and other extracellular matrix proteins, consequently destroying the hematopoietic microenvironment. A pivotal role for certain pro-fibrotic and pro-inflammatory growth factors, such as megakaryocyte-derived transforming growth factor β (TGF), is well recognized (*7–9*). However, the complexity of cell lineages that send and receive the signals that fuel bone marrow inflammation and fibrosis has not been fully elucidated. For example, while various mesenchymal stromal cell (MSC) subsets have been studied, including Nestin+ (*10*), Gli1+ (*11*) and Leptin-receptor+ MSCs (*12*), little is known about the subtypes and transcriptional states of bone marrow fibroblasts in myelofibrosis, and the specific cellular mediators and receptor-ligand (R-L) interactions that lead to pathological stromal cell activation.

Our aim was to build a comprehensive atlas of myelofibrotic bone marrow including hematopoietic stem/progenitor cells (HSPCs), mature hematopoietic cells and their stromal cell neighbors, to identify potentially targetable mediators of inflammation and fibrosis. To achieve this, we first mapped the cellular and molecular cross-talk in myelofibrotic bone marrow at single cell resolution in a mouse model of myelofibrosis, and corroborated findings using bone marrow biopsies and blood samples from patients. Unexpectedly, we found that basophils and mast cells – populations not previously highlighted as important inflammatory drivers in MPNs – are increased in abundance and act as the ‘hub’ for the enhanced TNF signaling. We also showed that while MSCs divert to produce extracellular matrix (ECM) components and downregulate their production of hematopoietic support factors, a compensatory increase in production of hematopoietic cytokines occurs from basophils, mast cells and a subset of pro-inflammatory bone marrow fibroblasts (iFibs). Therefore, paracrine hematopoietic support within myelofibrotic bone marrow derives from alternative cellular sources to that in healthy marrow.

The ý-galactoside binding protein galectin 1 emerged as one of only two genes differentially expressed in both the MPN clone and the inflamed stroma in myelofibrosis. We confirmed a clear positive correlation between galectin 1 expression and myeloid cancer progression in three large patient cohorts, and showed that inhibition of galectin 1 using a neutralizing anti-galectin 1 monoclonal antibody (mAb) ameliorated myeloproliferation and fibrosis in a mouse model and in 3D, multi-lineage human bone marrow organoids. This identifies galectin 1 as a robust biomarker and a therapeutic target for MPNs and potentially other myeloid malignancies and fibrotic disorders.

## RESULTS

### Generating a high-resolution cellular atlas of myelofibrotic bone marrow

To enable detailed analysis of the cellular landscape of myelofibrotic bone marrow, we utilized a well-characterized murine model in which MPL^W515L^, the third most common driver mutation occurring in MPN patients, is introduced into murine HSPCs by retroviral transduction and the cells transplanted into lethally irradiated, wild-type recipients (*13*). Control mice received HSPCs transduced with GFP alone. As previously described, this resulted in a severe and rapidly progressive myeloproliferative disease that was typically lethal within 4 weeks (*13*). Mice receiving MPL^W515L^ bone marrow developed leukocytosis, thrombocytosis, polycythemia, pronounced splenomegaly, a reduction in body weight, bone marrow fibrosis, reduced cellularity and increased, atypical megakaryocytes (Figure 1A and B, Supplemental Figure 1A). Histology of the spleens revealed loss of the normal lymphoid follicle architecture and markedly increased splenic megakaryocytes and other myeloid cells (Supplemental Figure 1B).

**Figure 1.**
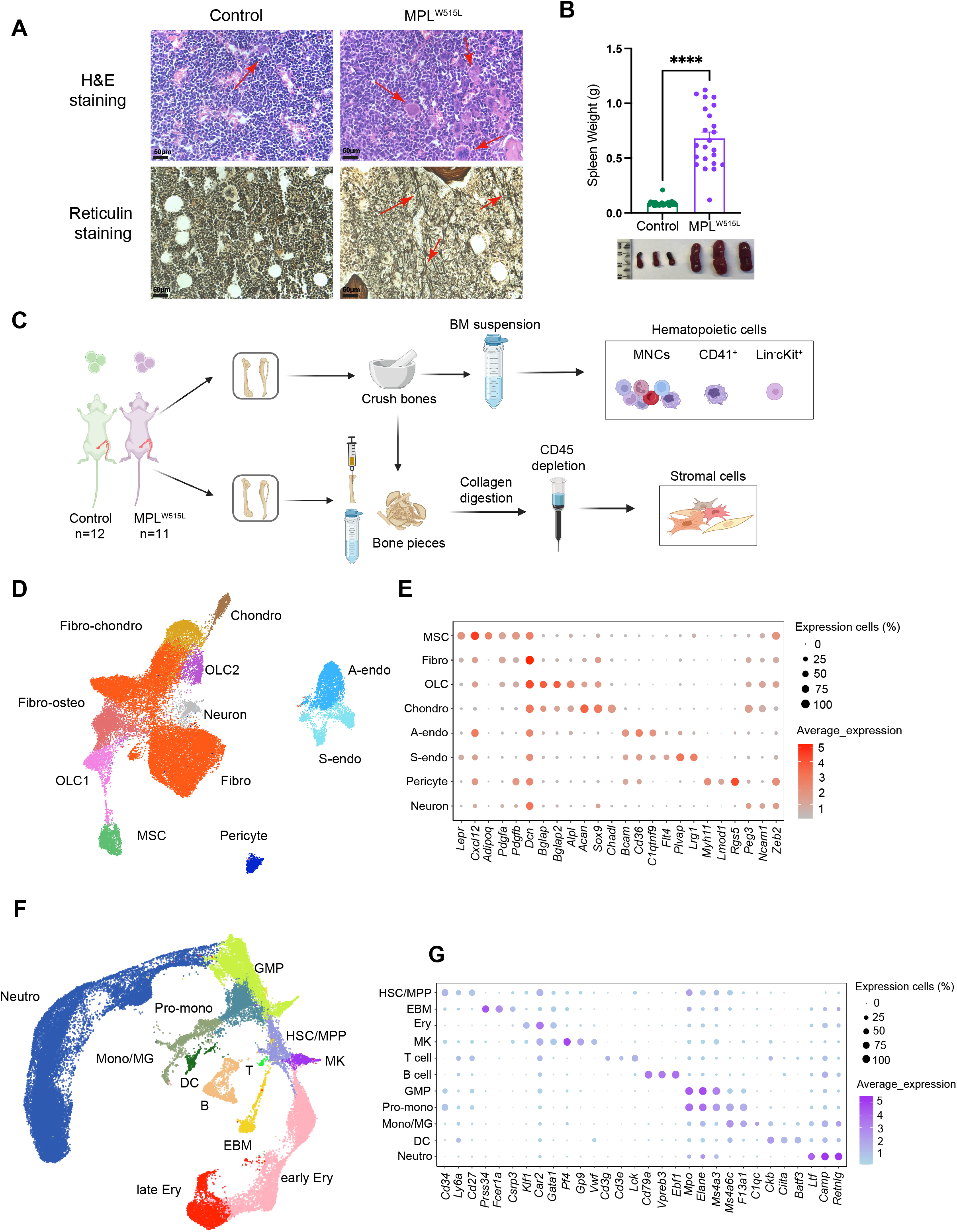
A high resolution cellular atlas of myelofibrotic bone marrow. (**A**) H&E (top) and reticulin stained (bottom) femur sections from control (n=9) and MPL^W515L^ mice (n=13). Red arrows highlight megakaryocytes (top) and reticulin fibrosis (bottom), representative images shown. (**B**) Spleen weights (grams, g) with representative images of control (n = 24) and MPL^W515L^ (n=24) mice. ****p < 0.0001 for unpaired t test with Welch’s correction. Chart shows mean ± SEM. (**C**) Schematic of experimental workflow to capture hematopoietic cells including lineage negative (lin-) cKit+ HPSCs, CD41+ and total mononuclear cells, as well as bone marrow stromal cells from control (n=12) and MPL^W515L^ mice (n=11). n=3 experiments. Created with Biorender.com. (**D and F**) Uniform Manifold Approximation and Projections (UMAPs) of (**D**) 34,969 stromal cells and (**F**) 42,319 hematopoietic cells from 12 GFP control mice and 11 MPL^W515L^ mice, colored by annotated cell cluster. (**E and G**) Dot plots showing expression of canonical marker genes used to annotate **(E)** stromal and (**G**) hematopoietic cells. Abbreviations: BM, bone marrow; MNC, mononuclear cells; Fibro-chondro, fibroblast-chondrocytes; Chondro, chondrocytes; OLC, osteolineage cells; Fibro-osteo, fibroblast-osteoblasts; Fibro, Fibroblasts; MSC, mesenchymal stromal cells; A-endo, arterial endothelial cells; S-endo, sinusoidal endothelial cells; Neutro, neutrophils; GMP, granulocyte-monocyte progenitors; Pro-mono, monocyte progenitors; Mono/MG, monocyte/macrophages; HSC/MPP, hematopoietic stem and multipotent progenitor cells; MK, megakaryocytes; EBM, eosinophil, basophil, mast cells; DC, dendritic cells; B, B-cell; T, T-cell; Ery, erythrocytes.

We devised a workflow enabling simultaneous capture of hematopoietic and stromal cells from murine femurs, tibiae and iliac crests, and performed high-throughput, droplet-based, single cell RNA-sequencing, isolating total mononuclear cells (MNCs) and enriching for rarer relevant cell types including lineage negative (Lin-) cKit+ HSPCs and cells expressing the megakaryocyte cell surface marker CD41 (Figure 1C). Capture of non-hematopoietic stromal cells was achieved by performing collagenase digestion of flushed and crushed bone pieces, beads depletion of CD45+ hematopoietic cells and then fluorescence-activated cell sorting (FACS) to isolate the CD45-, Lin-, Ter119-, CD71mid/-non-hematopoietic cell fraction (Figure 1C, Supplemental Figure 1C and Supplemental Table 1) (*14*).

Following data integration, doublet removal and quality control (Supplemental Figure 1D), 77 288 cells were analyzed from 23 mice in 3 independent experiments, including 42 319 hematopoietic and 34 969 stromal cells, generating a comprehensive atlas of normal and myelofibrotic bone marrow (Figure 1D – 1G, dataset available via an online data explorer at http://16.16.110.183:3838/myelofibrosisatlas/). Differentially expressed genes for each cluster were calculated after dimensional reduction and clustering, and cell types identified by their expression of canonical marker genes (Figure 1E and 1G, Supplemental Figure 1E, Supplemental Table 2).

We successfully captured the major cellular subsets annotated in recently published atlases of murine bone marrow (*14–17*). Within the bone marrow stroma, this included: MSCs (expressing *Lepr*, *Cxcl12*, *Adipoq*), fibroblasts (*Dcn*, *Pdgfra*, *Pdgfrb*) (*18*), osteolineage cells (OLC, *Bglap*, *Bglap2*, *Alpl*), chondrocytes (*Acan*, *Sox9*), pericytes (*Myh11*, *Rgs5*) (*19*) and neuronal (*Ncam1*) cells, and distinct arteriolar (*Bcam*, *C1qtnf9*) and sinusoidal (*Plvap*, *Lrg1*) endothelial cell subtypes (Figure 1D and 1E, Supplemental Figure 1E). Eleven hematopoietic cell lineages were captured, including hematopoietic stem and multipotent progenitor cells (HSC/MPP, *Cd34*, *Ly6a*, *Cd27*), megakaryocytes (*Pf4*), T (*Lck*) and B (*Cd79a*, *Ebf1*, *Vpreb3*) lymphocytes, eosinophil/basophil/mast cells (*Prss34*, *Fcer1a*), erythroid (*Car2*, *Gata1*), granulocyte-monocyte progenitors and pro-monocytes (*Mpo*, *Elane*), monocytes/macrophages (*Ms4a6c*) and neutrophils (*Camp*, *Retnlg*; Figure 1F & 1G, Supplemental Figure 1E).

To compare the cell types captured in our study to previously published datasets of normal (*14*) and myelofibrotic (*20*) bone marrow, Symphony (*21*) analysis was performed, using our data as the reference dataset and projecting cells from existing datasets onto to the reference embeddings. This confirmed annotation in our dataset of several major cell types including fibroblasts, chondrocytes, endothelial, osteolineage, mature neutrophils, eosinophils, basophils, and mast cells that were not captured in previous studies of myelofibrotic bone marrow, particularly in the stromal cell compartment (Supplemental Figure 2A and 2B) (*17, 20*). This dataset therefore represents an unbiased cellular and molecular atlas of the bone marrow in myelofibrosis, enabling a more comprehensive analysis of cellular and molecular interactions and perturbations than has been possible to date.

**Figure 2.**
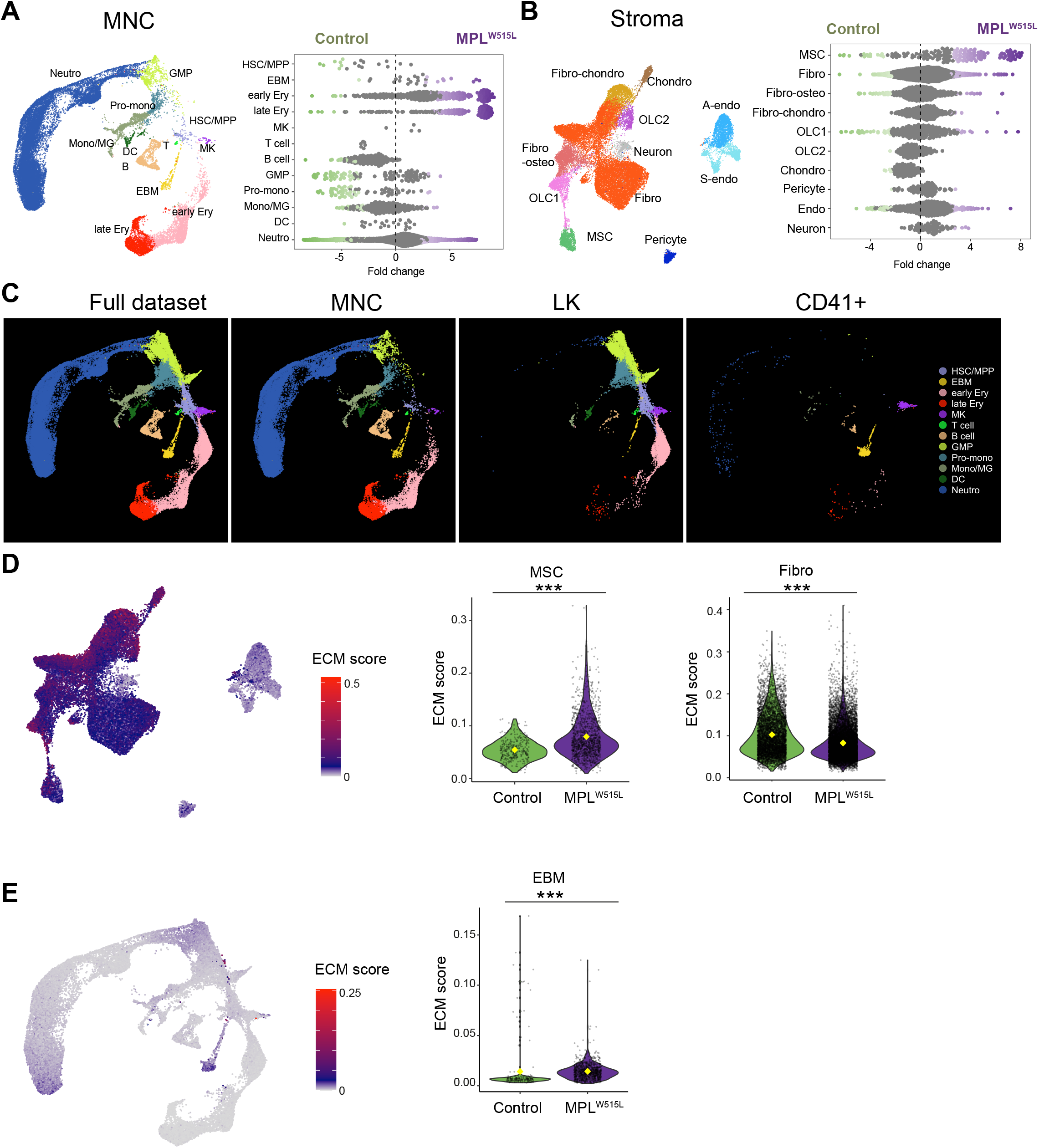
Alterations to the cellular constituents of myelofibrotic bone marrow and source of extracellular matrix components. **(A and B)** Differential abundance of **(A)** mononuclear cell (MNC) subsets and (**B**) stromal cell subsets in control (green) vs. MPL^W515L^ mice (purple), shown with relevant Uniform Manifold Approximation and Projections (UMAPs) to indicate relative frequency of each cell type. Each dot in the differential abundance plots represents a KNN cluster of the indicated cell type, clusters marked green and purple indicate those significantly depleted or enriched in MPL^W515L^ mice respectively. Sinusoidal and arterial endothelial cells are merged (endo) for the purpose of differential abundance in panel B. (**C**) Derivation of the total bone marrow hematopoietic cells captured (full dataset) from the three flow cytometric sorting strategies for MNCs, lineage negative cKit+ HSPCs (LK) and CD41+ cells (CD41), indicating that eosinophil, basophil and mast (EBM) cells and megakaryocytes (MK) were primarily captured by the CD41+ cell sort. (**D and E**) UMAPs (left) showing expression of a gene set of extracellular matrix factors (ECM) in (**D**) stromal and (**E**) full hematopoietic cell dataset, with violin plots (right) showing expression in relevant cell clusters from control (green) and MPL^W515L^ mice (purple). Yellow diamond indicates mean value. ***p < 0.001 for Wilcoxon test.

### Alterations to the cellular constituents of the bone marrow in myelofibrosis

The relative abundance of cell lineages was substantially altered in myelofibrotic bone marrow. In concordance with the expected disease phenotype, erythroid, neutrophil and megakaryocyte cells were substantially expanded in the hematopoietic compartment of MPL^W515L^ mice (Figure 2A). A pronounced decrease in HSPCs was also observed, with a near absence of B and T lymphocytes (Figure 2A). Within the stromal compartment, the most striking change in myelofibrosis mice was an expansion of MSCs, with a more modest increase in fibroblasts and decrease in chondrocytes, OLCs and endothelial cells compared to controls (Figure 2B).

Notably, basophils and mast cells were markedly increased in abundance in myelofibrosis mice compared to controls (Figure 2A). Unexpectedly, we found that the eosinophil, basophil and mast cell (EBM) population had primarily been captured by the enrichment sort for CD41+ cells (Figure 2C), a canonical cell surface marker of megakaryocyte cells but also expressed on murine basophils at steady-stage, and upregulated after cytokine activation (*22*).

### Altered cellular sources of ECM components

A defining feature of myelofibrosis is the aberrant deposition of ECM in the bone marrow, causing reticulin fibrosis, bone marrow failure and extramedullary hematopoiesis in particular in the spleen. The specific constituents and cellular origin of ECM factors in normal and myelofibrotic bone marrow have not been well described, although ECM components are recognized as important regulators of HSC function (*23*). To determine the cellular sources of ECM proteins in the bone marrow, we utilized an ECM gene list derived from proteomic analysis of normal and malignant tissues (*24*). Higher numbers of ECM genes were expressed by cells from the stroma than the hematopoietic compartment (total ECM genes: n = 233 *vs*. 107; collagens: n = 142 *vs*. 7; glycoproteins: n =159 *vs*. 76 and proteoglycans: n = 32 *vs*. 14 for stroma vs. hematopoietic respectively). Within the stromal cell subsets, high per cell expression of ECM genes was detected in all cell types apart from endothelial cells, neurons and pericytes (Figure 2D, Supplemental Figure 2C). Expression of collagen subtypes and glycoproteins was higher in OLCs and chondrocytes than other stromal cell subtypes, while fibroblasts and fibro-chondrocytes were the primary cellular source of proteoglycans, and MSCs predominantly expressed glycoproteins (Supplemental Figure 2C).

Expression of ECM components were also detected in the hematopoietic compartment, although in lower abundance than in the stroma (Figure 2E). Notably, prominent expression of glycoproteins and proteoglycans were detected in EBM cells, as well as a small fraction of monocytes/macrophages and mature neutrophils (Supplemental Figure 2D).

In myelofibrotic bone marrow, *per cell* expression of ECM genes was substantially increased in MSCs and EBM cells but decreased in fibroblasts (Figure 2D and 2E), suggesting that MSCs and EBM cells are major contributors to the altered deposition of extracellular matrix proteins in myelofibrosis.

### Altered cellular sources of hematopoietic support factors in myelofibrosis

Bone marrow Lepr+ MSCs transdifferentiate into myofibroblasts in myelofibrosis in response to platelet derived growth factor receptor (PDGFR) stimulation, downregulating their production of hematopoietic niche support factors in parallel with their increased expression of fibrogenic and osteogenic genes (*12, 20*). We detected clear transcriptional reprogramming of MSCs in myelofibrotic bone marrow, with a pronounced reduction in expression of hematopoietic niche support factors (Figure 3A, Supplemental Table 4) in parallel with the increased expression of ECM factors (Figure 2D). While the reduction in expression of hematopoietic support factors by MSCs in myelofibrosis has been documented (*12, 20, 25*), prior studies did not examine whether the production of hematopoietic support ‘shifts’ from MSCs to other cellular components of the bone marrow niche. We found that the reduction in supportive cytokines from MSCs was compensated by a significant increase in hematopoietic support from fibroblasts and also EBM cells in myelofibrosis *vs*. control cells (Figs. 3A and 3B). The *per cell* expression of niche support factors, in particular *Cxcl12* and *Csf1*, was markedly decreased in myelofibrosis vs. control bone marrow MSCs but increased in fibroblasts (Supplemental Figure 3A and B).

**Figure 3.**
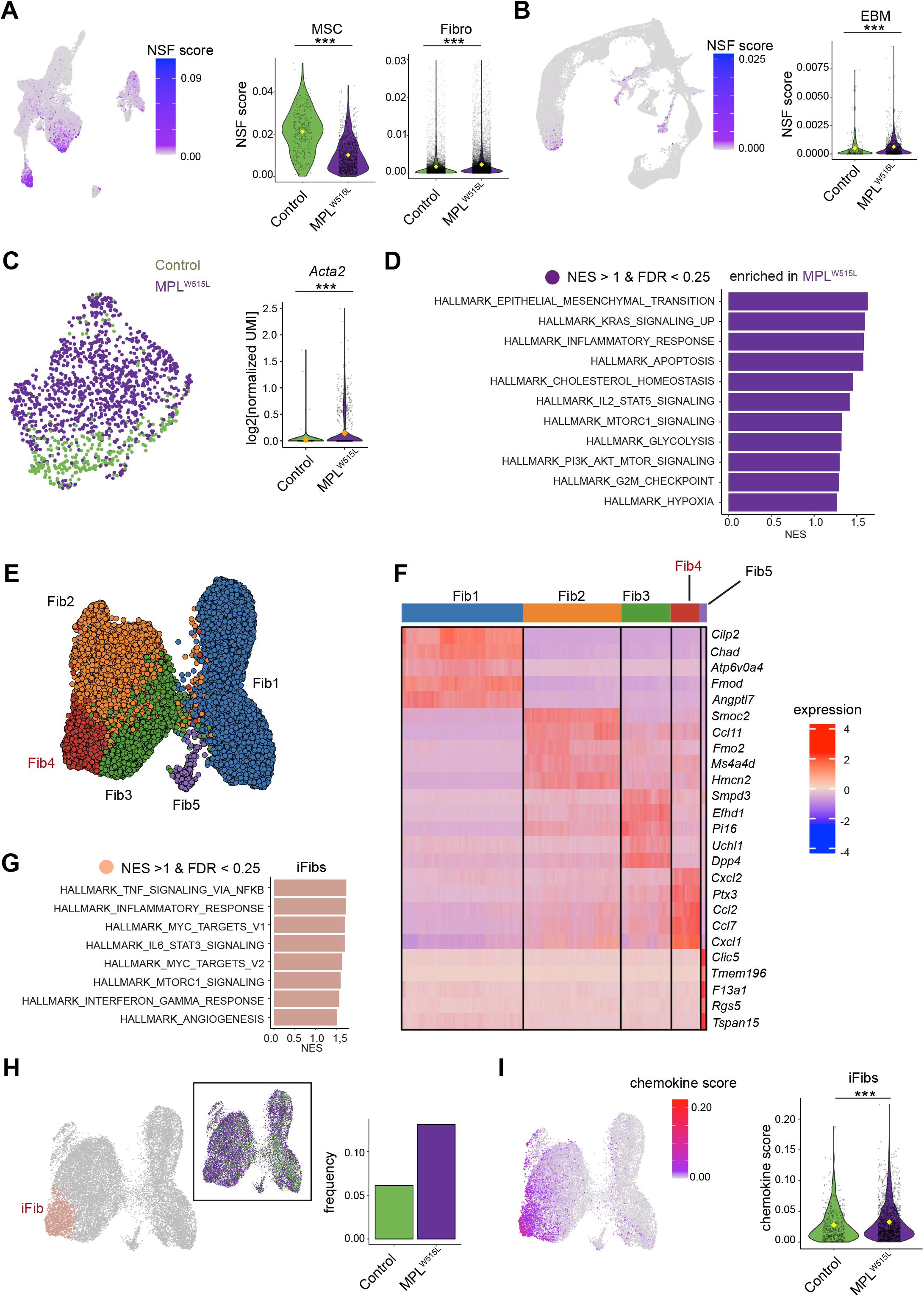
Altered cellular sources of hematopoietic support factors and expansion of inflammatory fibroblasts in myelofibrosis bone marrow. (**A & B**) Uniform manifold Approximation and Projection (UMAP, left) and violin plots (right) showing expression of niche supporting factors (NSFs) in (**A**) stromal and (**B**) hematopoietic cell datasets. Violin plots show expression in mesenchymal stromal cells (MSC), fibroblasts (Fibro) and eosinophil, basophil & mast cells (EBM) from control (green) and MPL^W515L^ mice (purple). ***p < 0.001 for Wilcoxon test. (**C**) MSCs from control (green) and MPL^W515L^ mice (purple) cluster separately, reflecting marked transcriptional reprogramming and myofibroblast trans-differentiation as indicated by increased alpha smooth muscle actin (*Acta2*). (**D**) Significantly enriched HALLMARK gene sets in MSCs from myelofibrosis mice. Selected gene sets shown. (**E**) UMAP showing 5 fibroblast sub-clusters. (**F**) Top 5 differentially expressed genes in each fibroblast subcluster. (**G**) Selected HALLMARK gene sets significantly enriched in Cluster 4, reflecting inflammatory fibroblast (iFib) phenotype. (**H**) Frequency of iFibs in MPL^W515L^ *vs*. control mice. (**I**) Expression of chemokine genes in fibroblasts. ***p < 0.001 for Wilcoxon test.

Myelofibrosis MSCs were transcriptionally distinct, with significant enrichment of pathways associated with myofibroblast transition, KRAS and phosphoinositide-3-kinase (PI3K) signaling, inflammatory response genes and IL2-STAT5 signaling (Figure 3C and 3D). Therefore, myelofibrosis-induced MSC trans-differentiation leads to increased ECM production but reduced hematopoietic support from MSCs, indicating that hematopoiesis is guided by alternative cellular sources in the setting of MPNs, potentially influencing the competitive advantage of the MPN clone over healthy hematopoiesis.

### Emergence of a distinct inflammatory fibroblast subset in the myelofibrotic niche

The relative proportion of fibroblast cells overall was only minimally increased in myelofibrotic bone marrow (Figure 2B). As fibroblasts were the most abundant stromal cell type captured, and as distinct fibroblasts subsets have been reported to be important in other pathologies (*26, 27*), we extracted the fibroblasts for further analysis, confirming their expression of the canonical fibroblast markers *Pdgfra/Pdgfrb* and performing unsupervised sub-clustering (Supplemental Figure 3C).

Five transcriptionally distinct sub-clusters were identified (Figure 3E, Supplemental Table 2), of which one cluster (Fib4) uniquely showed striking enrichment for inflammatory pathways (Figure 3F and 3G) and was therefore annotated as representing inflammatory fibroblasts (iFibs). iFibs were significantly enriched for TNF signaling via NFKB, inflammatory response signaling, IL6-JAK-STAT3 signaling and interferon gamma response (Figure 3G). The relative frequency of iFibs was 2-fold higher in myelofibrosis mice than controls (Figure 3H), and expression of chemokine genes was strongly enriched in the iFibs with significantly increased per cell expression of chemokines in MPL vs. control fibroblasts (Figure 3I, Supplemental Table 4), including *Kitl*, *Cxcl12*, *Ccl2*, *Cxcl1* (Supplemental Figure 3A & Supplemental Figure 3D and 3E). The iFib cluster also expresses *Cxcl5*, which has been identified in a recent fibroblast atlas as a marker for perturbation-specific, activated fibroblast states and not detected in steady-state fibroblasts (*18*) (Supplemental Figure 3F). Collectively these data support that, although overall fibroblast numbers are only slightly altered in myelofibrosis, distinct inflammatory fibroblast subsets producing hematopoietic support factors are markedly expanded in number, thereby contributing to the development of an aberrant hematopoietic niche in myelofibrosis.

### Expanded pro-inflammatory basophils, mast cells and megakaryocytes in myelofibrosis

Megakaryocyte proliferation and morphological atypia are hallmark features of overt and pre-fibrotic myelofibrosis (*1*), and we found an expansion of megakaryocytes with angiogenic, proliferative and inflammatory gene expression programs in the myelofibrosis mice (MK3, 4 and 5, Supplemental Figure 4A – E). While megakaryocytes are well recognized as important drivers of fibrosis (*9, 28*), the pathological contributions of basophil and mast cell subsets in myelofibrosis have not been extensively studied (*29*). Having noted a significant increase in the abundance of EBM cells (Figure 2A), we extracted cells from the EBM cluster for a more detailed analysis (Figure 4A). Four distinct subtypes of EBM cells were annotated – EBM progenitors, mast cells, basophils and a small population of mature eosinophils (Figs. 4A and 4B, Supplemental Figure 4F, Supplemental Table 2). The relative proportions and transcriptional activity of these cellular subsets were very distinct in myelofibrosis bone marrow, with a dramatic expansion of basophils and mast cells, and relatively few eosinophils in myelofibrosis mice compared to controls (Figure 4C), and significant enrichment of IL2-STAT5, TGFB, and TNF via NF-κΒ inflammatory signaling pathways (Figure 4D and 4E).

**Figure 4.**
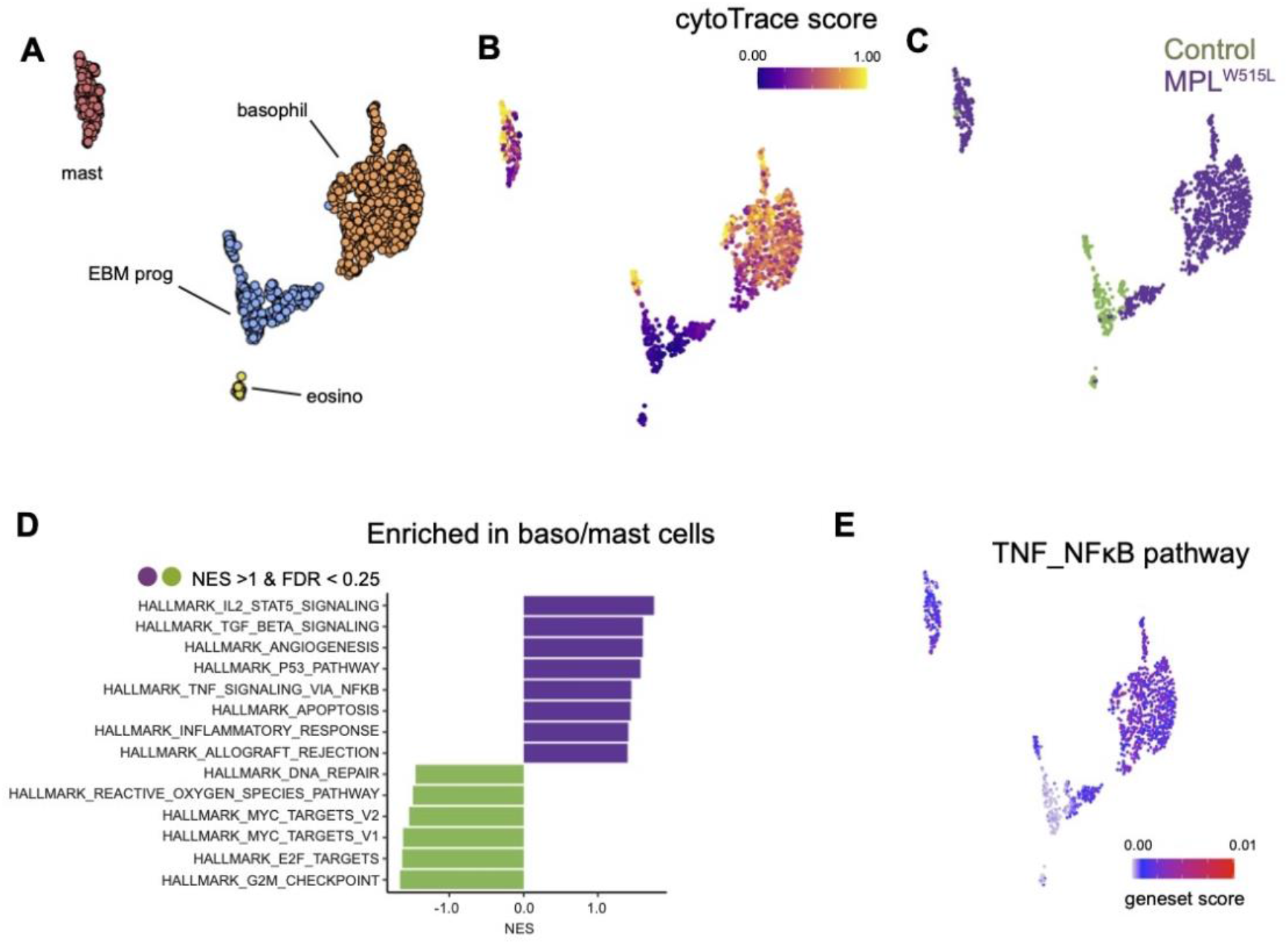
Expansion of pro-inflammatory basophils and mast cells in myelofibrosis. (**A**) UMAP showing annotated sub-clusters of cells from the eosinophil, basophil and mast (EBM) cell cluster. (**B**) CytoTRACE differentiation state analysis of EBM cells, with blue indicating primitive state and yellow showing differentiation trajectory. (**C**) UMAP identifying cells originating from MPL^W515L^ (purple) and control (green) mice. (**D**) Significantly enriched HALLMARK gene sets in basophils and mast cells from MPL^W515L^ vs. control mice. (**E**) Expression of TNF-NFκB pathway genes projected onto the EBM cell UMAP.

### Basophils and mast cells emerge as the ‘hub’ of TNF and pro-inflammatory cytokine signaling

To identify how the cellular cross-talk was altered in myelofibrotic bone marrow, we computationally inferred the interacting receptor-ligand (R-L) pairs that might mediate communication between cell types (*30*). The overall number of predicted R-L interactions was 20% higher in MPL^W515L^ than control mice (Figure 5A), and the aberrant signaling was largely due to increased interactions deriving from basophils, mast cells and megakaryocytes in the hematopoietic compartment and MSCs and inflammatory fibroblasts in the stroma (Figure 5B), highlighting these 4 cell types as ‘orchestrators’ of inflammatory signaling in myelofibrotic bone marrow. Basophils and mast cells emerged as the hub of TNF and IL4 signaling in MPL^W515L^ mice, with fibroblasts, inflammatory fibroblasts, MSCs and neutrophils and monocytes/macrophages as their key interacting partners (Figure 5C and 5D, Supplemental Figure 5A & 5B).

**Figure 5.**
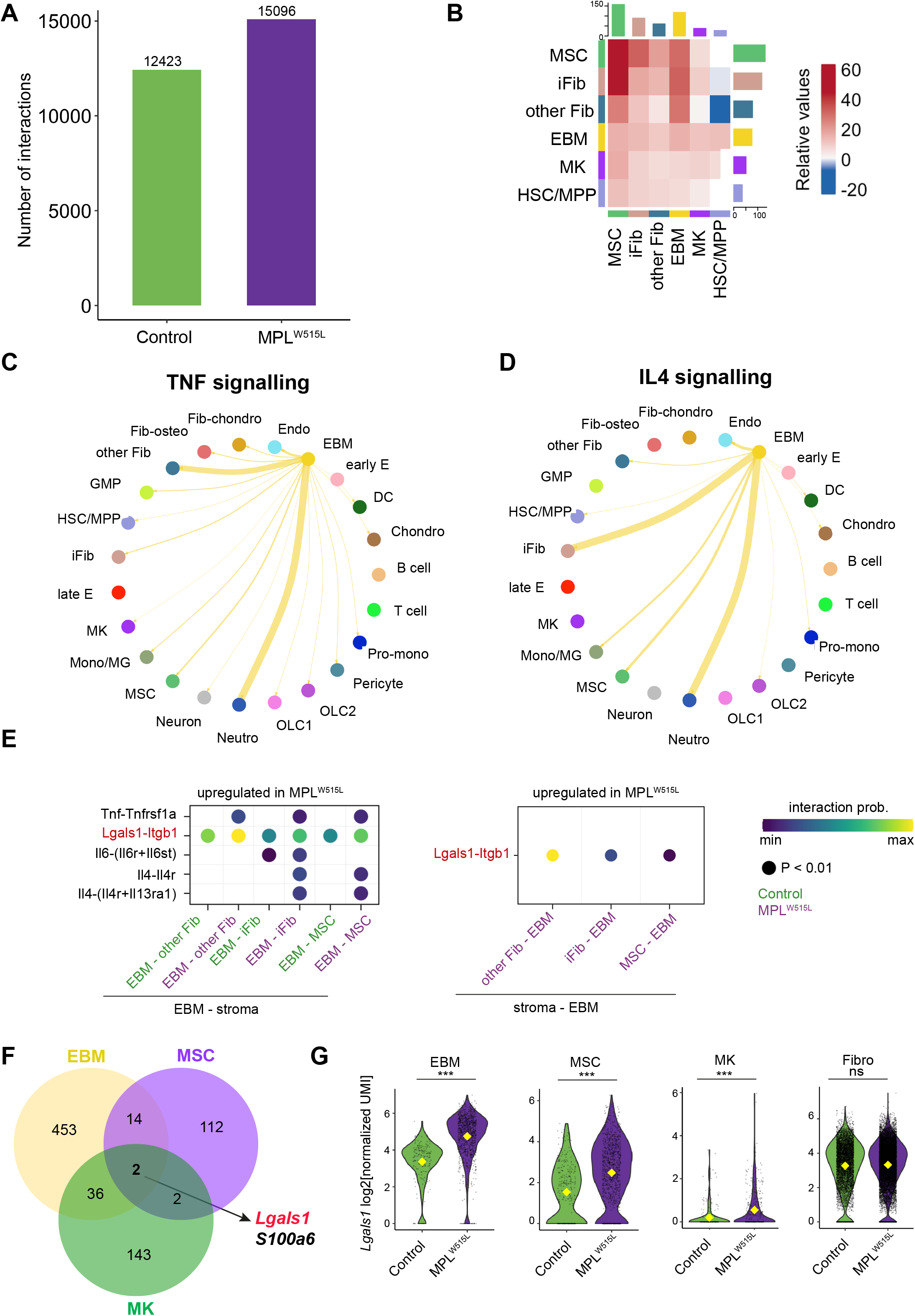
Basophils and mast cells emerge as the ‘hub’ of TNF and interleukin 4 pro-inflammatory signaling. (**A**) Number of inferred Ligand (L) – Receptor (R) interactions in control and MPL^W515L^ bone marrow. (**B**) Differential number of L-R interactions in MPL^W515L^ vs. control bone marrow. Total Number of enriched L-R interactions is shown as a bar on the x/y axes and relative strength of the interactions (MPL^W515L^ vs. control bone marrow) is shown in the heatmap for key stromal and hematopoietic cell populations. (**C & D**) Circus plot depicting interaction pathway of (**C**) TNF and (**D**) IL4 uniquely upregulated in MPL^W515L^ mice. The width of the connections reflects the strength of the interactions between two populations. (**E**) Selected L-R interactions predicted to be upregulated in MPL^W515L^ mice between EBM cells, fibroblasts, iFibs, and mesenchymal stromal cells (MSCs). (**F**) Venn diagram showing distinct and overlapping differentially expressed genes in EBM, MSC and MK clusters. (**G**) Violin plots showing expression of *Lgals1* in EBM, MSC, MK and fibroblasts in control and MPL^W515L^ mice. Abbreviations: R – L, receptor-ligand; L – R, ligand-receptor; TNF, tumor necrosis factor alpha; IL, interleukin; EBM, eosinophil, basophil, mast cells; iFibs, inflammatory fibroblasts; MSCs, mesenchymal stromal cells; Fibro, fibroblast; HSC/MPP, hematopoietic stem and multipotent progenitor cells. ***p < 0.001; ns – non-significant for Wilcoxon test.

*Lgals1*, the gene encoding the protein galectin 1, a ý-galactoside binding protein which interacts with ý-1 integrin (*Itgb1*), emerged as a R-L pair with substantially enhanced predicted signaling across the key interacting cell types (Figure 5E, Supplemental Figs. 5C and 5D). When we looked for genes which were differentially expressed in the key interacting cell types in myelofibrosis, only two genes were concordantly dysregulated across cell types – *S100a6* and *Lgals1* (Figure 5F). A role for *S100a6* and other *S100* family members in inflammation and malignant hematopoiesis has previously been reported (*31–33*), whereas galectin 1 has not been extensively studied in myeloid malignancies. Expression of *Lgals1* was strikingly increased in basophils and mast cells, MSCs and megakaryocytes in MPL^W515L^ mice compared to control mice, with high levels of expression in fibroblasts overall but no significant difference in per cell expression level (Figure 5G). Together, these data suggested that galectin 1 signaling might play a key pathological role in myelofibrosis progression and warranted further exploration.

### Galectin 1 inhibition ameliorates myelofibrosis disease phenotype *in vivo*

To test whether galectin 1 signaling contributes to the pathobiology of myelofibrosis *in vivo*, we tested the impact of a neutralizing anti-galectin monoclonal antibody (Gal-1-mAb3) that binds to a specific sequence in galectin 1 not present in other galectin family proteins (*34*) in the MPL^W515L^ mouse model. Control and MPL^W515L^ mice were treated with either IgG isotype control or Gal-1-mAb3 by intraperitoneal injection (Figure 6A). Galectin 1 neutralization led to a reduction in bone marrow fibrosis and cellular architecture in the MPL^W515L^ mice (Figure 6B), and reduced the myeloproliferative phenotype with significantly reduced thrombocytosis, polycythemia and splenomegaly (Figure 6C – E). Notably, the reduction in splenomegaly with Gal-1-mAb3 treatment was similar to that with fedratinib, a JAK2 inhibitor in clinical use, in this model (Figure 6F), and no cytopenias were observed following galectin 1 inhibition in the control mice (Figure 6C & 6D), indicating specific inhibition of the MPN clone rather than a non-specific cytoreductive impact. Furthermore, inhibition of galectin 1 led to significantly improved MPN-free survival (Supplemental Figure 6A).

**Figure 6.**
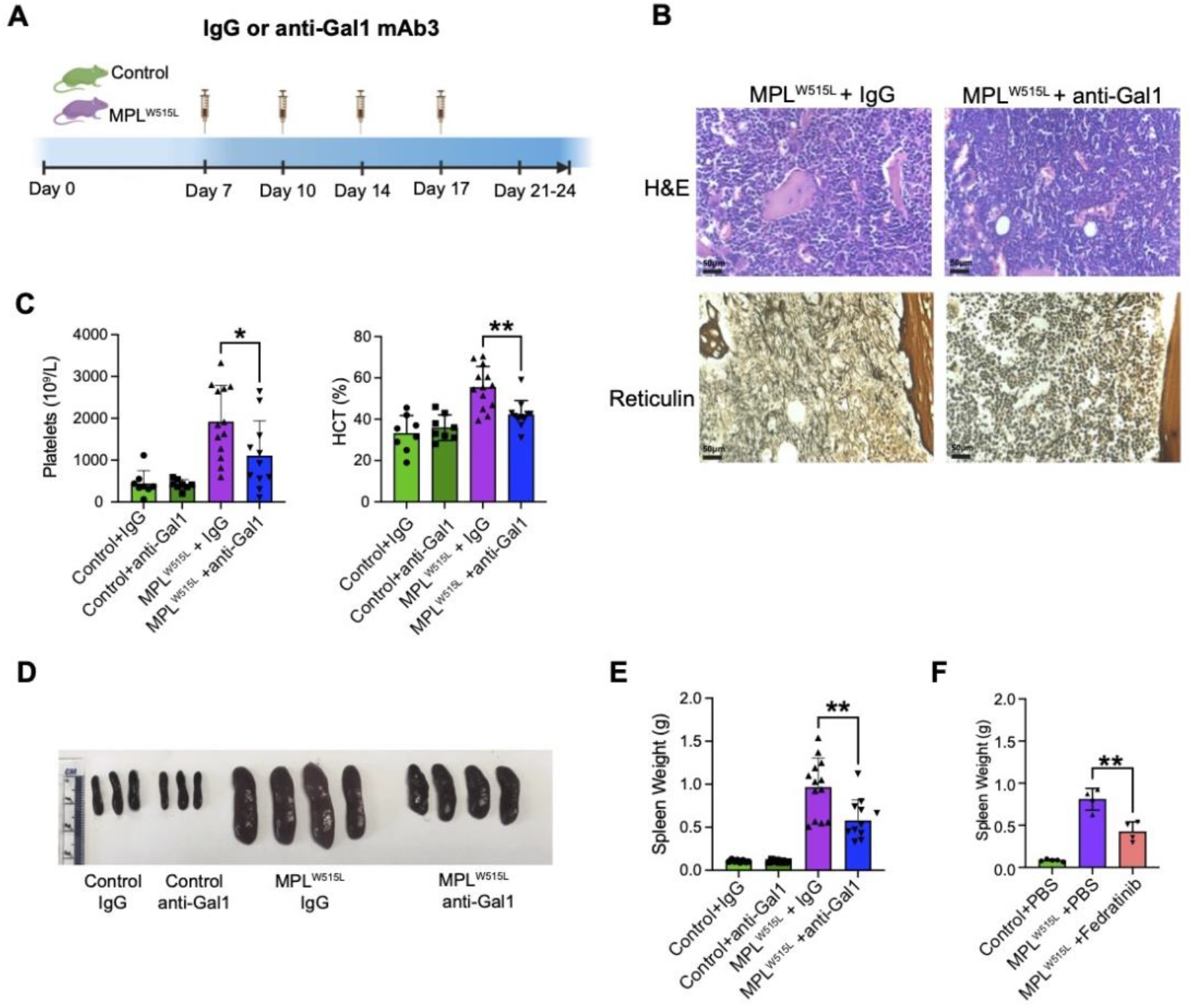
Inhibition of Galectin 1 ameliorates fibrosis and myeloproliferation in vivo. **A**) Schematic of treatment with isotype control (IgG) or anti-Gal1 mAb3, initiated on day 7 following transplantation of control or MPL^W515L^ BM cells. (**B**) H&E and reticulin staining of femur sections from MPL^W515L^ mice treated with IgG control or anti-Gal1 mAb3. Representative images shown. (**C**) Platelet counts and hematocrit (HCT) in IgG or anti-Gal1 treated control (n=8 and n=8) and MPL^W515L^ mice (n=13 and n=11). *p < 0.05, **p < 0.01 for unpaired t test with Welch’s correction. Data represented as mean ± SEM. (**D & E**) Representative images (**D**) and weights (**E**) of spleens from IgG or anti-Gal1 mAb3 treated control (n=8 and n=8) and MPL^W515L^ mice (n=13 and n=11). (**F**) Mean ± SEM spleen weights of mice treated with PBS control (n=5) or the JAK2 inhibitor fedratinib (n=4). *p < 0.05, **p < 0.01 for unpaired t test with Welch’s correction.

### Galectin 1 is a robust biomarker of fibrosis progression in patients with MPNs

Given the amelioration of disease phenotype *in vivo* in the mouse model, we next sought to validate galectin 1 in myeloid malignancies in the setting of human disease, using a series of patient cohorts. We first tested whether galectin 1 expression correlated with fibrosis progression in patients with myeloproliferative neoplasms, quantifying galectin 1 protein in bone marrow biopsies of 30 patients, including those with myelofibrosis (n = 14), non-fibrotic MPNs (essential thrombocythemia [ET], n = 9 and polycythemia vera [PV], n = 7) and age-matched healthy controls (n = 7) (Supplemental Table 3). Galectin 1 was markedly increased in myelofibrotic bone marrow (Figure 7A). Objective quantification of staining intensity per high power field view showed a significant increase in galectin 1 with progression to myelofibrosis across patient groups (Figure 7B, P < 0.001 for myelofibrosis vs. healthy donors and P < 0.001 for myelofibrosis vs. ET and PV). Bone marrow fibrosis is often unevenly distributed in the bone marrow space, and this heterogeneity is inadequately captured by the standard categorical fibrosis grading system that is typically employed in clinical assessments (e.g. WHO grade MF 0 – 3). In order to measure the association between galectin 1 expression and reticulin fibrosis more precisely, we employed a recently developed machine learning pipeline that enables automated fibrosis quantification by allocating a Continuous Index of Fibrosis (CIF) score for each bone marrow region, creating a heatmap representing the density of fibrosis across the entire marrow specimen (*35*). This showed clear correlation between the intensity of galectin 1 immunostaining and the density of fibrosis within the marrow sections (Figure 7C), as well as between patient samples (Figure 7B).

**Figure 7.**
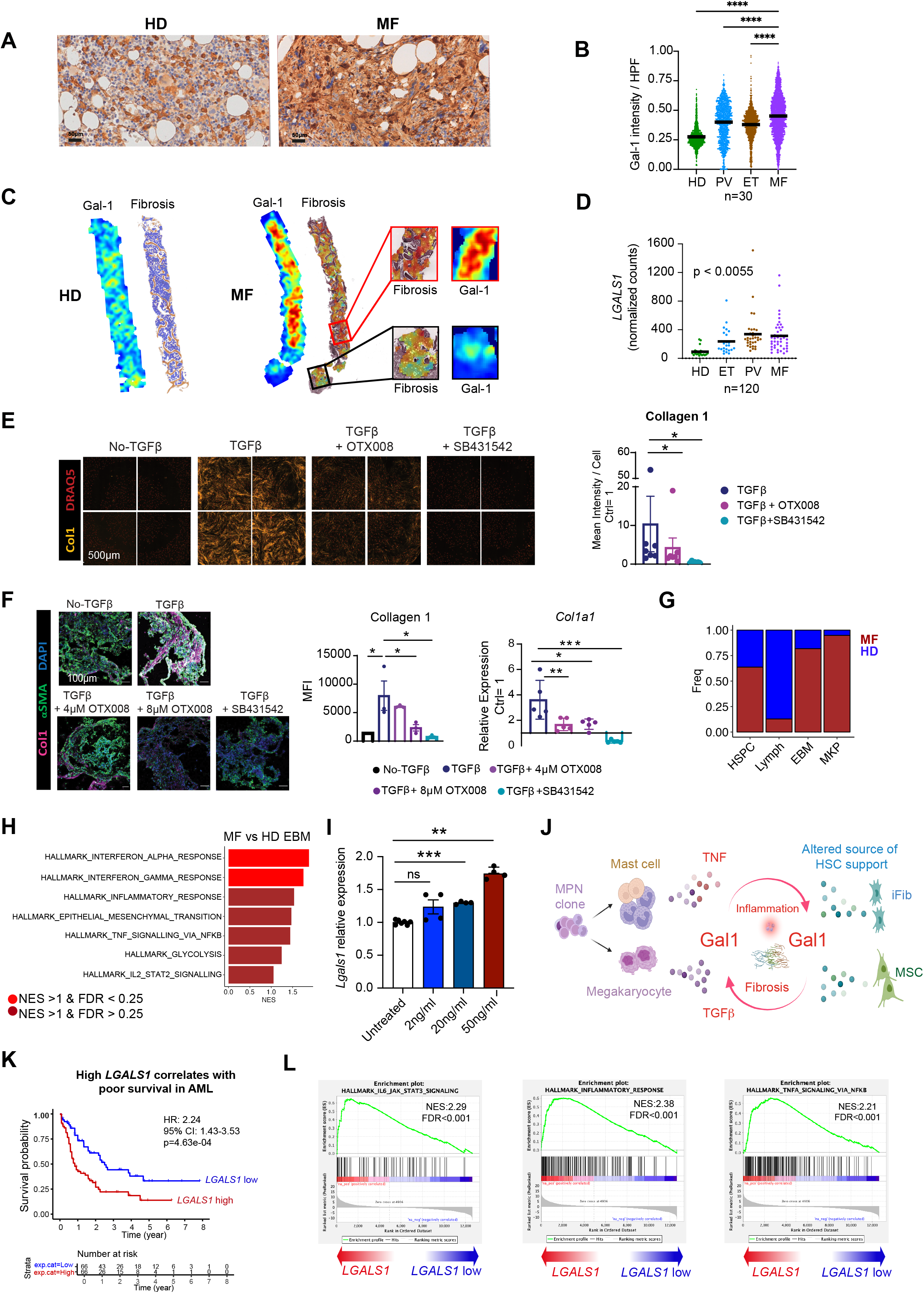
Galectin 1 is a robust biomarker for fibrosis and poor outcomes in myeloid malignancies. **(A)** Representative immunohistochemistry staining for galectin 1 (Gal1) in healthy donor (HD) (n=7) and myelofibrosis (MF) (n=14) bone marrow biopsy sections. Data are represented as mean ± SEM. **(B)** Gal1 expression per high power field (HPF) view in bone marrow biopsy sections from HDs (n=7) and patients with essential thrombocythemia (ET, n = 9), polycythemia vera (PV, n=7), and MF (n=14). ****p < 0.0001 for Kruskal-Wallis test. Data are represented as mean ± SEM. **(C)** Gal1 staining intensity correlated with reticulin fibrosis density across bone marrow biopsy sections from HDs and MF patients. Color scale from blue to red as fibrosis density increases. Representative images shown. **(D)** LGALS1 expression in platelets from a cohort of 120 HDs and patients with MPNs (HD=21, ET=24, PV=33, MF=42). Data are represented as mean ± SEM. (**E**) TGFβ-induced fibroblast to myofibroblast transition assay using human BMSCs treated with TGFβ alone ± OTX001 (galectin 1 inhibitor) or SB431542 (TGFβ inhibitor). Representative images shown for high-throughput, 384-well imaging plate (left). Each treatment was performed in quadruplicate and 4 images acquired per well (n=7 patients). Chart (right) shows MFI per cell for collagen 1 normalized to the no-TGFβ control ± SEM (n=7). *p < 0.05 for wilcoxon matched pairs signed rank test. (**F**) Impact of OTX008 on TGFβ-induced Collagen 1 and aSMA in human iPSC-derived BM organoids. Representative images (left); Mean ± SEM for protein/mRNA expression quantification of Collagen 1/*COL1A1* (right). n=5-8 organoids from 3 independent experiments. *p < 0.05, **p < 0.01, ****p < 0.0001 for one-way ANOVA. (**G**) Bar chart showing relative proportion of cell subtypes from a previously published dataset of ∼120,000 cells from human CD34+ hematopoietic stem/progenitor cells from patients with myelofibrosis (MF) and age-matched healthy donors (HD). (**H**) Enriched HALLMARK gene sets in EBM progenitor cells from MF patients *vs*. HD. Abbreviations: UMAP, Uniform manifold Approximation and Projection; EBM, eosinophil (eosino)-basophil (baso)-mast cells; MF, myelofibrosis; HD, healthy donors; NES, normalized enrichment score; FDR, false discovery rate. **(I)** *Lgals1* mRNA expression in human bone marrow organoids with/without treatment with TNF at doses shown. Data are represented as mean ± SEM, n=80 organoids across 2 independent experiments. *p < 0.05, **p < 0.01, ****p < 0.0001 for one-way ANOVA. (**J**) Schematic illustrating the interactions between basophils, mast cells and megakaryocytes derived from the MPN clone interacting with BMSC subsets, fueling inflammation and fibrosis via galectin 1 induction. Created with Biorender.com. Abbreviations: Ctrl, control; TGFβ, transforming growth factor β; Col1, collagen 1; αSMA, alpha smooth muscle actin; anti-Gal1, monoclonal anti-Galectin 1 neutralizing antibody; HCT, hematocrit; IgG, isotype IgG control antibody; H&E, hematoxylin and eosin; g, grams. (**K**) Kaplan-Meier survival curves showing correlation between high *LGALS1* expression and poor survival in 132 patients with acute myeloid leukemia (AML) in The Cancer Genome Atlas (TCGA) dataset. (**L**) Gene set enrichment analysis show significant enrichment of IL6-JAK-STAT3 signalling, inflammatory response and TNF signalling via NFKB in patients with high *LGALS1* expression in TCGA database. Abbreviations: HR, hazard ratio; NES, normalized enrichment score; FDR, false discovery rate.

To further validate galectin 1 as a biomarker and to see if it could be utilized as a non-invasive peripheral blood biomarker of fibrosis, we investigated galectin 1 expression in a large cohort of 120 patients where platelet transcriptomes were available from patients with myelofibrosis (n=42), ET (n=24), PV (n=33) and healthy controls (n=21) (*36*). A progressive and highly significant increase in galectin 1 expression was observed with progression of MPN to fibrosis (monotonic trend from controls to PV/ET to myelofibrosis, P < 0.0055, Figure 7D), with a 3.4-fold increase in myelofibrosis vs. controls.

### Galectin-1 validates as a targetable mediator of fibrosis in human cellular assays and bone marrow organoids

MPN mouse models are useful surrogates for the human disease, but evidence that a potential target can be functionally validated using human experimental systems is more compelling. We therefore explored whether galectin 1 was mediating a severe disease phenotype using human disease models. We derived bone marrow stromal cells (BMSCs) from marrow aspirates of patients with MPNs (Supplemental Table 3) and utilized these in a TGFβ-induced fibroblast-to-myofibroblast transition assay (*37*). Treatment of BMSCs with recombinant human TGFβ led to increased collagen 1 deposition and αSMA expression, which was reversible on inhibition of TGFβ signaling with SB431542, an inhibitor of the TGFβ activin receptor-like kinase (ALK) receptors (*38*). OTX008, a small molecule galectin 1 inhibitor previously shown to inhibit pulmonary fibrosis, inhibited TGFβ-induced fibroblast-to-myofibroblast transition (Figure 7E and Supplemental Figure 6B) (*39*).

To confirm a role for galectin 1 as a mediator of TGFý-induced bone marrow fibrosis in a multi-cellular bone marrow microenvironment, we utilized a three dimensional model that better recapitulates the complexity of human bone marrow. Bone marrow organoids were generated from human induced pluripotent stem cells using an optimized protocol that gives rise to the key stromal and hematopoietic cellular elements of the central marrow space, approximating the transcriptional and architectural features of the native human hematopoietic tissues (*40*). In this model, OTX008 significantly inhibited TGFý-induced collagen 1 and αSMA expression at both protein and mRNA level (Figure 7F and Supplemental Figure 6C).

### TNF upregulates galectin 1 gene expression

Given the pronounced increase in TNF signaling from basophils and mast cells in the MPL^W515L^ mouse model (Figures 4E and 5C), and as TNF-NF-κý signaling has previously been shown to regulate LGALS1 expression by T-cells(*41*), we hypothesized that TNF might stimulate galectin 1 production in human bone marrow. We first corroborated that basophils and mast cells were increased in frequency and had an inflammatory phenotype in the setting of myelofibrosis in patients by interrogating a scRNAseq dataset of ∼120,000 CD34+ Lin-HSPCs isolated from a cohort of 15 myelofibrosis patients and 6 age-matched healthy donors (9). A population of EBM progenitors was identified (Supplemental Figure 6D, Supplemental Table 4), which were significantly more abundant in myelofibrosis patients than healthy controls (Figure 7G). Similar to our findings in the mouse model, these cells showed a striking enrichment of inflammatory response, IL2-STAT and TNF signaling (Figure 7H), confirming that basophils and mast cells are likely to play an important role in the pathobiology of myelofibrosis in patients and contribute to TNF pro-inflammatory pathways.

TNF is a potent activator of nuclear factor (NF)-κΒ(*42*), and NF-κΒ directly binds to regulatory elements in exon 1 of the *LGALS1* gene, enhancing protein expression(*41*). We therefore tested whether the mechanism of galectin 1 increase in myelofibrosis might occur secondary to TNF stimulation. Indeed, TNF treatment of bone marrow organoids robustly led to a three-fold increase in *LGALS1* expression (Figure 7I), suggesting a model wherein a self-reinforcing, inflammatory MPN niche is created by expanded populations of basophils, mast cells, MSCs and inflammatory fibroblasts with a central role for TGFý, TNF and galectin 1 signaling (Figure 7J).

### High galectin 1 predicts poor survival in acute myeloid leukemia

Given the disease modifying activity of the anti-galectin 1 antibody treatment in the MPL^W515L^ mouse model, we hypothesized that high expression of galectin 1 may be detrimental more broadly in myeloid malignancies. We therefore interrogated The Cancer Genome Atlas (TCGA) to test whether expression of galectin 1 correlated with overall survival in 132 patients with acute myeloid leukemia (*43*). There was a clear correlation between *LGALS1* expression level and poor survival (Figure 7K, P < 0.0005), with highly significant enrichment of inflammatory signaling pathways in patients with high *LGALS1* levels and poor survival, including inflammatory response, IL6 – JAK – STAT signaling and TNF signaling (Figure 7L).

Collectively, these results highlight galectin 1 as a central pathological mediator in myeloid malignancies, a promising biomarker, and a therapeutic target that may alter the disease course, which is not possible to achieve for the majority of patients using currently available medical therapies.

## DISCUSSION

MPNs are inflammatory pathologies that result in a significant burden of morbidity and mortality. The majority of patients present with early-stage malignancies, presenting an opportunity for intervention. However at present, there are no drug therapies that robustly impede or reverse progression to fibrosis, and a more detailed understanding of the genetic and non-genetic drivers of MPN progression is crucial. In this study, we present a comprehensive road-map of the cellular composition of myelofibrotic bone marrow, providing a platform for the discovery and characterization of novel cellular and molecular targets for therapy. Although prior studies highlighted important aspects of disease pathophysiology (*9, 17, 20*), these datasets have not simultaneously captured hematopoietic and stromal cells, precluding accurate delineation of the multi-lineage interactions that occur between myeloid cells of the MPN clone and components of their niche. The analyses presented here revealed perturbations to cellular frequencies and transcriptional phenotypes that were previously unappreciated, noting that an expansion of basophils, mast cells and a distinct subset of inflammatory fibroblasts collectively underlie pathogenic cellular interactions in myelofibrosis.

In individuals who acquire an MPN cancer driver mutation, the inflammatory microenvironment is an important determinant of clinical phenotype, symptom severity and the risk of disease progression (*44*). The same mutations can present with diverse clinical phenotypes, including in healthy individuals without overt hematologic disease (*45*). Although specific genetic contexts (high molecular risk mutations e.g., concurrent *ASXL1*, *SRSF2* mutations or a high *JAK2V617F* allele burden) increase the likelihood of progression to fibrosis, these are not essential, suggesting a major role for cell-extrinsic signaling in driving disease evolution. Recent studies revealed that MPN driver mutations are typically acquired early in life, often several decades before clinical presentation (*46–48*), yet myelofibrosis usually presents in the later decades of life. One explanation for the long latency observed between mutation acquisition and clinically overt disease is that the composition and function of the bone marrow stroma becomes more permissive for MPN outgrowth with age. A pro-inflammatory, TGFβ-rich stroma (*49*) and reduced MSC-derived hematopoietic support factors (*50*) develop with physiological ageing and induce a myeloid bias even in individuals without an MPN driver mutation. Here, we show that an MPN induces an exacerbation of the inflammatory and myeloid-biased hematopoiesis phenotype that occurs as part of healthy ageing (*51*), encouraging speculation that aging might accelerate the development of the self-reinforcing, malignant niche in myelofibrosis (*52*).

We demonstrate the utility of the dataset in identifying clinically-actionable targets by focusing on galectin 1, a β-galactoside binding protein that has been previously implicated in cancer, tissue fibrosis and immunoregulation (*53, 54*) but not myeloid malignancies. Exploration of galectin 1 expression levels in large patient cohorts showed a clear association with fibrosis progression and significant correlation with survival in patients with myeloid leukemias. A functional role for galectin 1 was confirmed, using 2D and 3D *in vitro* models of bone marrow fibrosis and also *in vivo* by demonstrating efficacy of a neutralizing anti-galectin 1 mAb (*34*).

Previous studies have suggested modes of action for galectin 1 that may be relevant in myeloid malignancies. Galectin 1 has been identified as a mediator of TGFβ- and hypoxia-induced lung fibrosis (*39*), and direct anti-proliferative effects have been shown using shRNA knock-down of galectin 1 as well as treatment with OTX008, a small molecule inhibitor that reached phase I clinical trials for patients with advanced solid tumors. Proliferative effects are mediated by ERK1/2 and AKT-dependent survival pathways, and galectin 1 inhibition induces of G2/M cell cycle arrest (*55*). Immunomodulatory activities are well documented for galectin 1, which acts as a suppressor of T cell anti-tumor immunity (*56*), enhances regulatory monocyte/macrophage subsets (*57*), promotes tolerogenic dendritic cells and in certain scenarios has been shown to trigger damage-associated molecular pattern (DAMP) pathway activation (*58*). Galectin 1 is a transcriptional target of NFκΒ, and its expression and release are enhanced via TNF signaling via NFκΒ (*59*). We show that a feedback loop exists wherein expanded basophil, mast cell, megakaryocyte and stromal cell subsets induce a self-reinforcing pro-inflammatory niche and galectin 1 expression, fueling inflammation and fibrosis (Figure 7J). Targeting galectin-1 using small molecule glycan inhibitors, natural polysaccharides, peptides (OTX008) or anti-galectin-1 monoclonal antibodies may counteract fibrosis and also the immunomodulation that occurs in myeloid neoplasms (*58, 60*).

Ongoing work will be aimed at determining the mechanisms of action for galectin 1 in myeloid neoplasms, further validating the efficacy of galectin 1 targeting in additional disease models, and identifying the most clinically-tractable targeting modality. Collectively, the data presented here confirm a role for galectin 1 as a mediator of pathobiology in myeloid malignancies and worthy of further exploration as a therapeutic target that has the potential to modify the disease course. The road-map of cellular interactions in myelofibrotic bone marrow has broad implications for other hematological malignancies, cancer-associated inflammation and non-malignant fibrotic disorders.

## One Sentence Summary

Unravelling the cellular landscape of myelofibrosis reveals novel drivers of inflammation and galectin 1 as a clinically actionable target.

## Acknowledgments

We thank Professor Ross Levine for sharing the MPL^W515L^ plasmid, the patients who kindly consented to research and: Patricia Ciccone, Nawshad Hayder and Sophie Reed who helped with sample banking; Kevin Clark, Craig Waugh and Paul Sopp in the MRC WIMM Flow Cytometry facility which is supported by the MRC Human Immunology Unit and MRC Molecular Haematology Unit; Dr. Neil Ashley in the MRC WIMM Single Cell Facility; Val Millar (Target Discovery Institute, University of Oxford); Ida Parisi (Histology Lab, Kennedy Institute); Ryan Beveridge (MRC WIMM Virus Screening Facility) and all the staff of the Biomedical Science Division. We thank Professor Jian Xu from University of Texas Southwestern Medical Center, for sharing the mouse HSPCs R objects (Liu et al, Nat Comms 2021).

## Author contributions

**Conceptualization, supervision and project administration:** B. Psaila, A.J Mead. **Funding acquisition:** B. Psaila, A.J Mead, R.Li. **Supervision of computational analysis:** G. Wang, S. Thongjuea. **Investigation:** R.Li, M. Colombo, A. Rodriguez-Romera, S. Clark, Y. Meng, A. O. Khan, L.C. Murphy, A.-A. Olijnik, Z.C. Wong, C. Simoglou Karali, R. Norfo, J. Carrelha, Z. Ren. **Validation:** R.Li, M. Colombo. **Methodology:** R.Li, M. Colombo, G. Wang, S.Wen, K. Sirinukunwattana, H. Ryou, Q. Cheng, A. Krishnan, D. Royston. **Original draft and editing:** R.Li, M. Colombo, G. Wang, B. Psaila, A.J Mead. **Resource**s: A. O. Khan, J. O’Sullivan, J.M. Pérez Sáez, N. Sousos, C.K. Brierley, V.A. Rathinam, D. Royston, G.A. Rabinovich. **Visualization:** R.Li, M. Colombo, G. Wang, P. Dong, W. Zhou, K. Sirinukunwattana, H. Ryou. **Data curation and analysis:** G. Wang, J. O’Sullivan, S.Wen, P. Dong, W. Zhou, K. Sirinukunwattana, H. Ryou, Q. Cheng, C.K. Brierley, S. Thongjuea, A. Krishnan. All authors read and approved the submitted manuscript.

## Data and materials availability

All raw and processed sequencing data generated in this study have been submitted to the NCBI Gene Expression Omnibus (GEO; https://www.ncbi.nlm.nih.gov/geo/) and the accession number will be made available upon formal publication. The corresponding sample information is contained in Suppl. Table 5. All code will be deposited at Github. Materials and reagents used in this study are listed in Supplementary Table 1.

