## Supplemental figures for "A pro-inflammatory stem cell niche drives myelofibrosis through a targetable galectin 1 axis"

### Figure S1

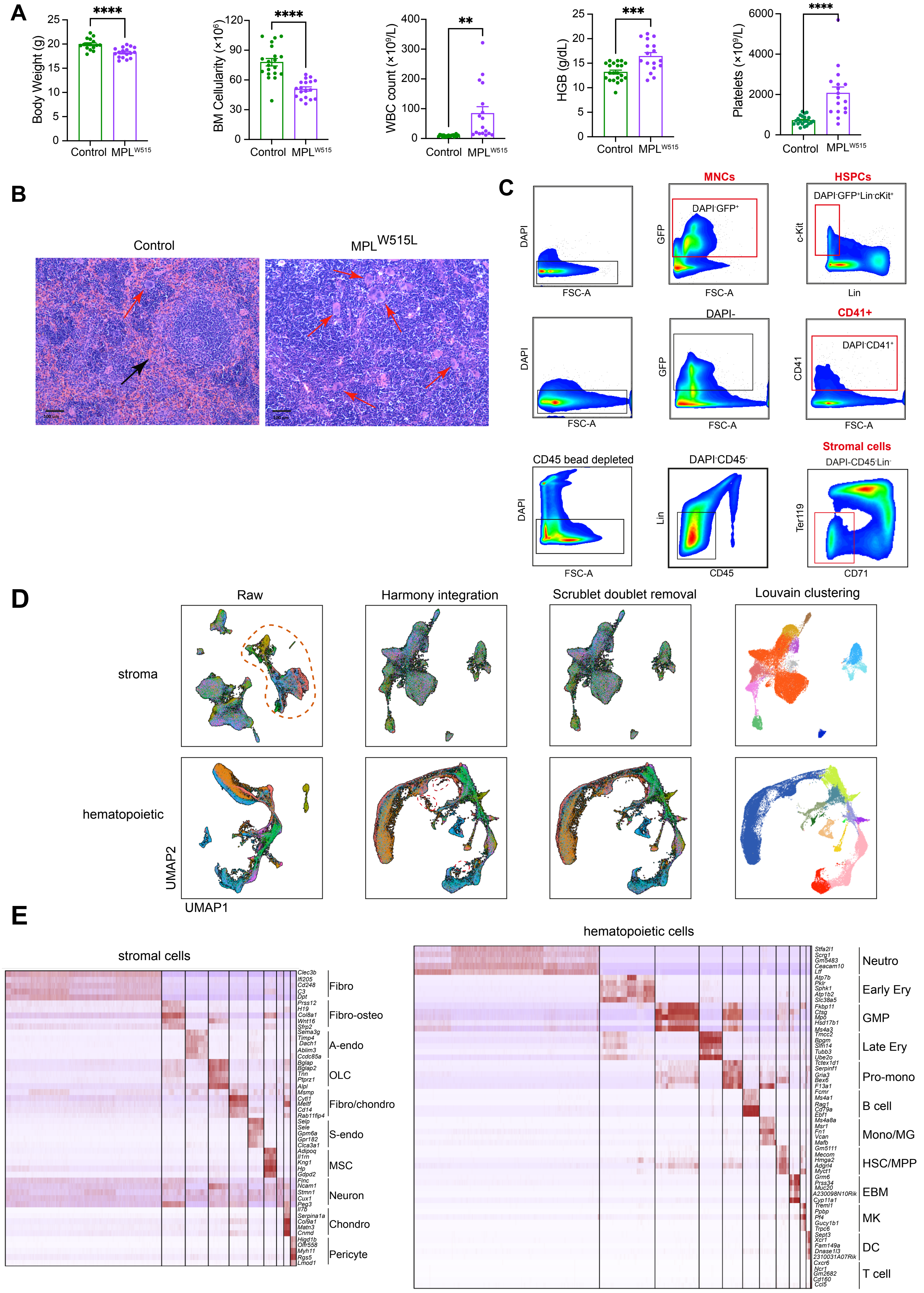

A
Body Weight (g)
****
Control
MPLW515
BM Cellularity (×10^6)
****
WBC count (×10^9/L)
**
HGB (g/dL)
***
Platelets (×10^9/L)
****
B
Control
MPLW515L
C
DAPI
FSC-A
MNCs
GFP
DAPI-GFP+
HSPCs
c-Kit
DAPI-GFP+Lin-cKit+
Lin
DAPI-
CD41+
CD41
DAPI-CD41+
CD45 bead depleted
DAPI-CD45-
CD45
Stromal cells
DAPI-CD45-Lin-
Ter119
CD71
D
Raw
Harmony integration
Scrublet doublet removal
Louvain clustering
stroma
hematopoietic
UMAP2
UMAP1
E
stromal cells
Fibro
Fibro-osteo
A-endo
OLC
Fibro/chondro
S-endo
MSC
Neuron
Chondro
Pericyte
hematopoietic cells
Neutro
Early Ery
GMP
Late Ery
Pro-mono
B cell
Mono/MG
HSC/MPP
EBM
MK
DC
T cell

### Figure S2

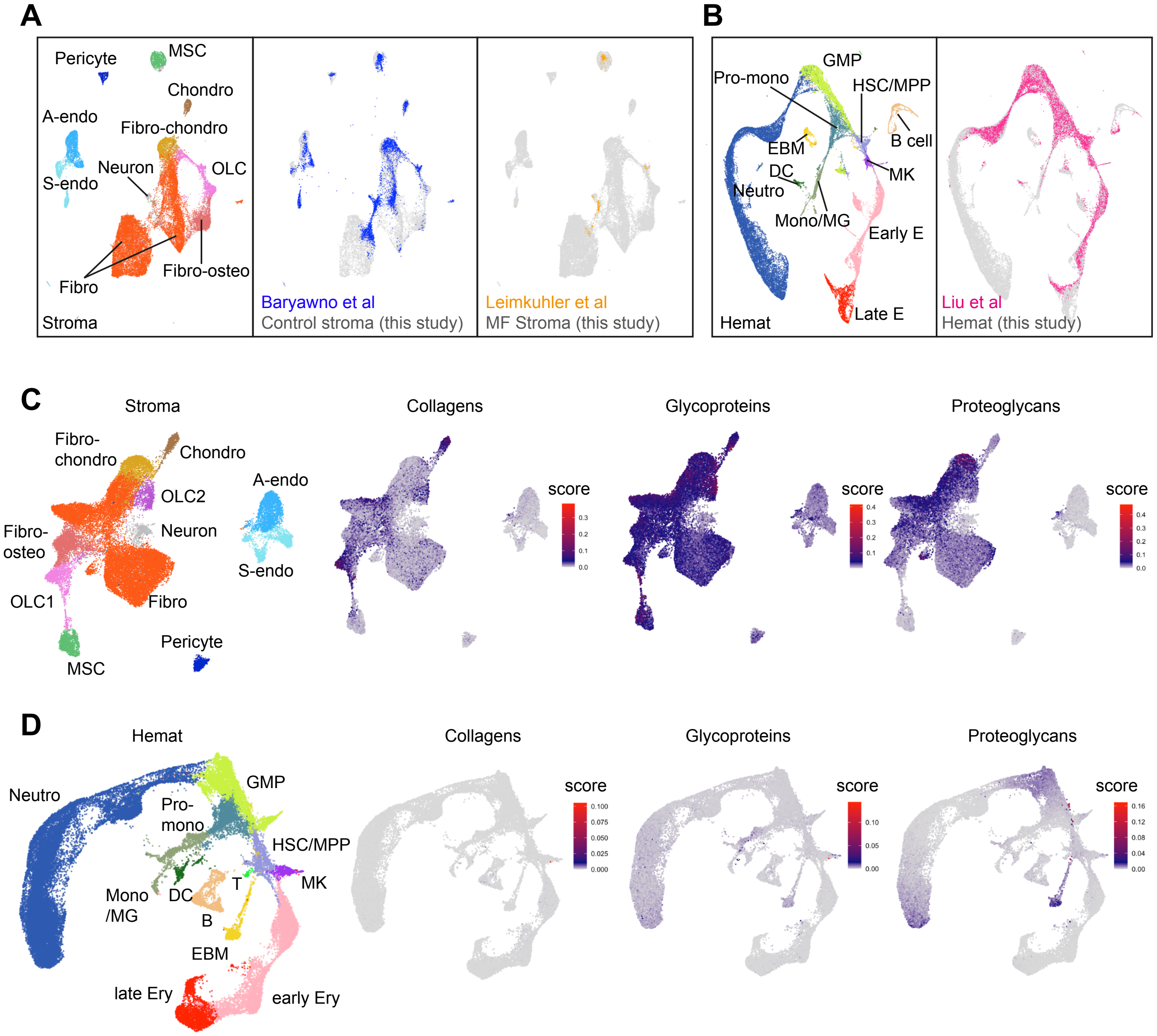

A
Pericyte
MSC
Chondro
A-endo
Fibro-chondro
Neuron
OLC
S-endo
Fibro-osteo
Fibro
Stroma
Baryawno et al
Control stroma (this study)
Leimkuhler et al
MF Stroma (this study)
B
GMP
Pro-mono
HSC/MPP
B cell
EBM
DC
MK
Neutro
Mono/MG
Early E
Late E
Hemat
Liu et al
Hemat (this study)
C
Stroma
Collagens
Glycoproteins
Proteoglycans
Fibro-
chondro
Chondro
A-endo
OLC2
Neuron
Fibro-
osteo
S-endo
OLC1
Fibro
Pericyte
MSC
score
D
Hemat
Collagens
Glycoproteins
Proteoglycans
Neutro
GMP
Pro-
mono
HSC/MPP
Mono
/MG
DC
T
MK
B
EBM
late Ery
early Ery
score

### Figure S3

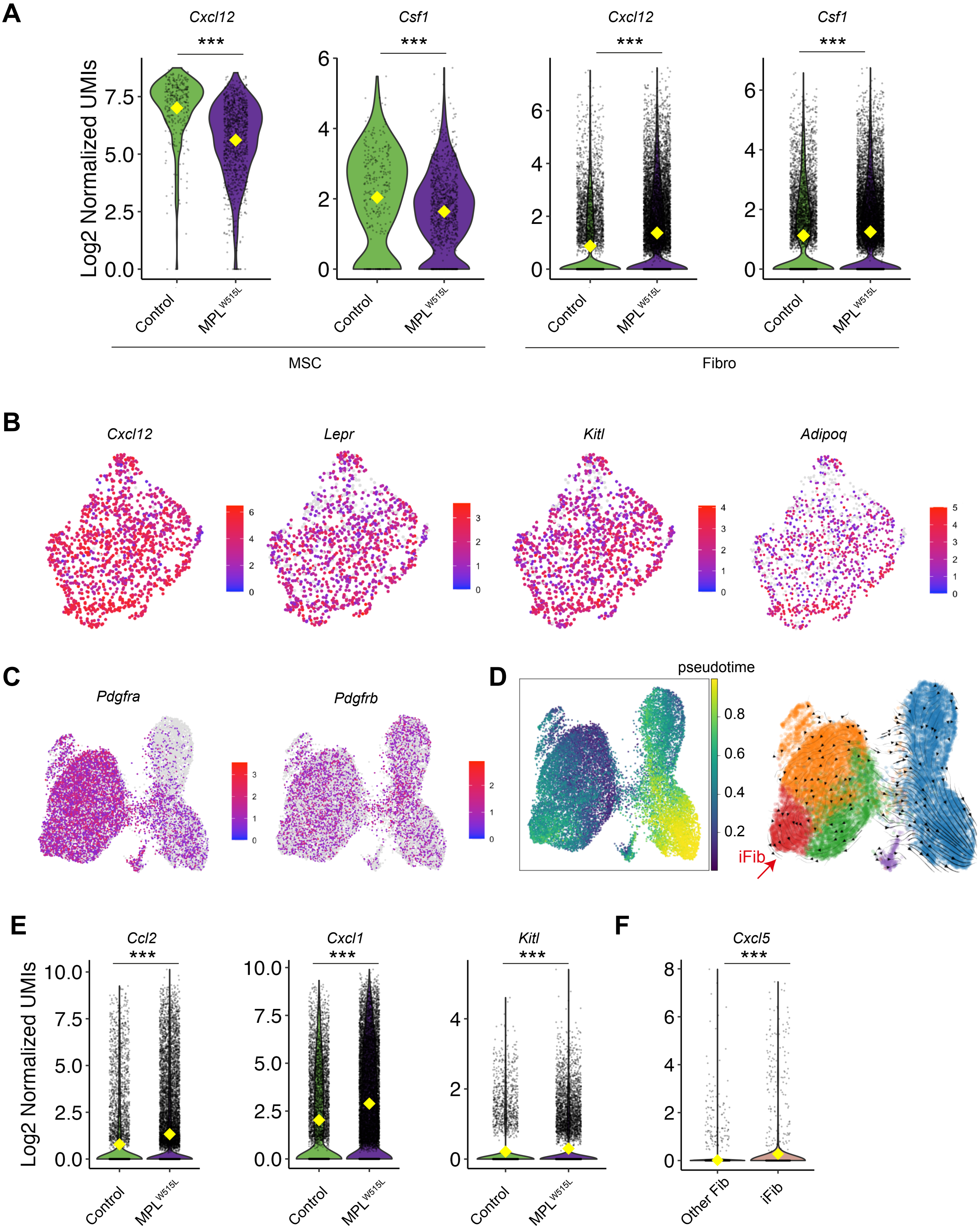
A
Cxcl12
***
Csf1
***
Cxcl12
***
Csf1
***
Log2 Normalized UMIs
Control
MPL W515L
MSC
Fibro
B
Cxcl12
Lepr
Kitl
Adipoq
C
Pdgfra
Pdgfrb
D
pseudotime
iFib
E
Ccl2
***
Cxcl1
***
Kitl
***
Log2 Normalized UMIs
Control
MPL W515L
F
Cxcl5
***
Other Fib
iFib

### Figure S4

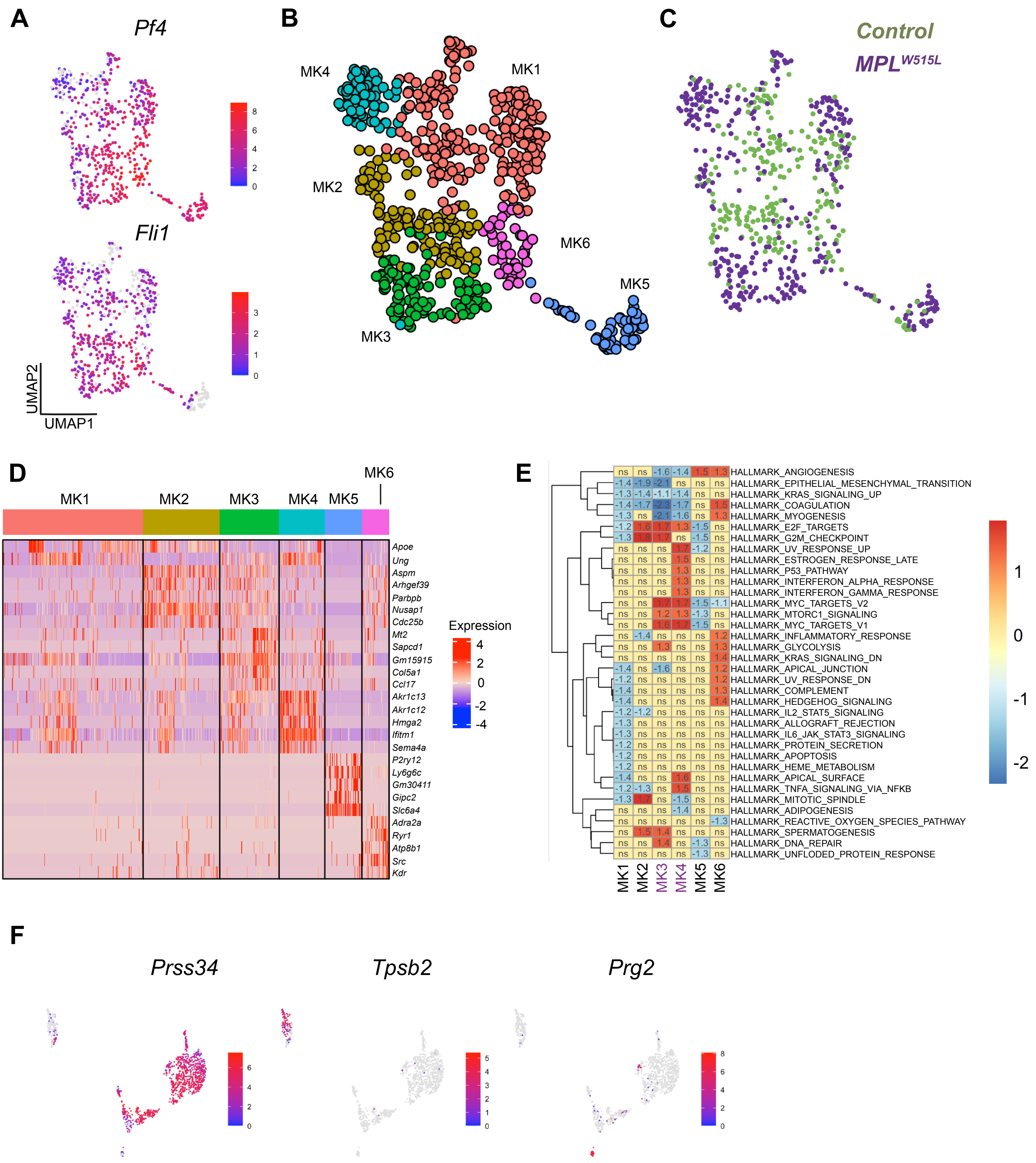
A
Pf4
Fli1
UMAP2
UMAP1
B
MK1
MK2
MK3
MK4
MK5
MK6
C
Control
MPL W515L
D
MK1
MK2
MK3
MK4
MK5
MK6
Apoe
Ung
Aspm
Arhgef39
Parbpb
Nusap1
Cdc25b
Mt2
Sapcd1
Gm15915
Col5a1
Ccl17
Akr1c13
Akr1c12
Hmga2
Ifitm1
Sema4a
P2ry12
Ly6g6c
Gm30411
Gipc2
Slc6a4
Adra2a
Ryr1
Atp8b1
Src
Kdr
Expression
E
HALLMARK_ANGIOGENESIS
HALLMARK_EPITHELIAL_MESENCHYMAL_TRANSITION
HALLMARK_KRAS_SIGNALING_UP
HALLMARK_COAGULATION
HALLMARK_MYOGENESIS
HALLMARK_E2F_TARGETS
HALLMARK_G2M_CHECKPOINT
HALLMARK_UV_RESPONSE_UP
HALLMARK_ESTROGEN_RESPONSE_LATE
HALLMARK_P53_PATHWAY
HALLMARK_INTERFERON_ALPHA_RESPONSE
HALLMARK_INTERFERON_GAMMA_RESPONSE
HALLMARK_MYC_TARGETS_V2
HALLMARK_MTORC1_SIGNALING
HALLMARK_MYC_TARGETS_V1
HALLMARK_INFLAMMATORY_RESPONSE
HALLMARK_GLYCOLYSIS
HALLMARK_KRAS_SIGNALING_DN
HALLMARK_APICAL_JUNCTION
HALLMARK_UV_RESPONSE_DN
HALLMARK_COMPLEMENT
HALLMARK_HEDGEHOG_SIGNALING
HALLMARK_IL2_STAT5_SIGNALING
HALLMARK_ALLOGRAFT_REJECTION
HALLMARK_IL6_JAK_STAT3_SIGNALING
HALLMARK_PROTEIN_SECRETION
HALLMARK_APOPTOSIS
HALLMARK_HEME_METABOLISM
HALLMARK_APICAL_SURFACE
HALLMARK_TNFA_SIGNALING_VIA_NFKB
HALLMARK_MITOTIC_SPINDLE
HALLMARK_ADIPOGENESIS
HALLMARK_REACTIVE_OXYGEN_SPECIES_PATHWAY
HALLMARK_SPERMATOGENESIS
HALLMARK_DNA_REPAIR
HALLMARK_UNFLODED_PROTEIN_RESPONSE
MK1
MK2
MK3
MK4
MK5
MK6
F
Prss34
Tpsb2
Prg2

### Figure S5

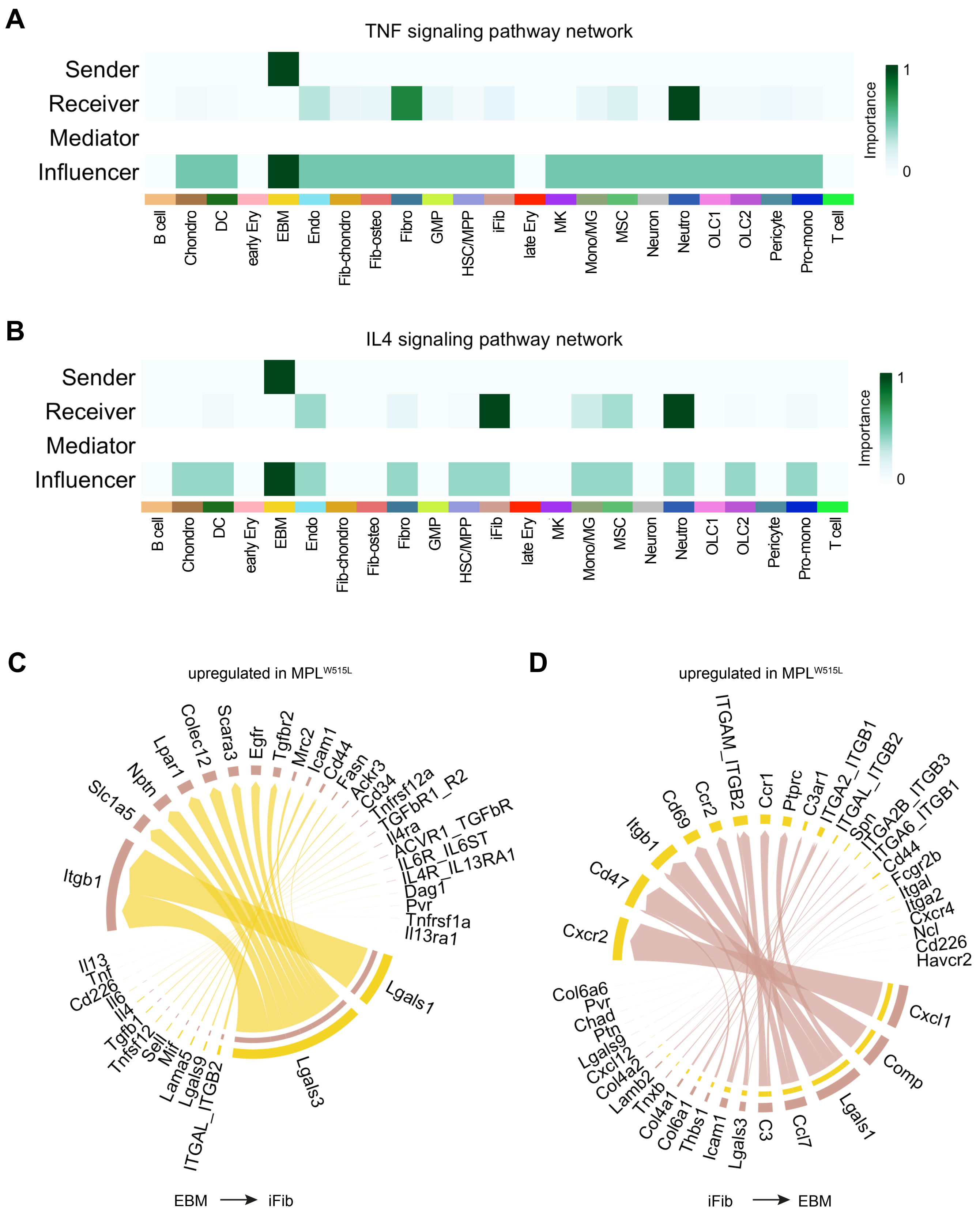

A
TNF signaling pathway network
Sender
Receiver
Mediator
Influencer
Importance
1
0
B cell
Chondro
DC
early Ery
EBM
Endo
Fib-chondro
Fib-osteo
Fibro
GMP
HSC/MPP
iFib
late Ery
MK
Mono/MG
MSC
Neuron
Neutro
OLC1
OLC2
Pericyte
Pro-mono
T cell
B
IL4 signaling pathway network
Sender
Receiver
Mediator
Influencer
Importance
1
0
B cell
Chondro
DC
early Ery
EBM
Endo
Fib-chondro
Fib-osteo
Fibro
GMP
HSC/MPP
iFib
late Ery
MK
Mono/MG
MSC
Neuron
Neutro
OLC1
OLC2
Pericyte
Pro-mono
T cell
C
upregulated in MPL W515L
Slc1a5
Nptn
Lpar1
Colec12
Scara3
Egfr
Tgfbr2
Mrc2
Icam1
Cd44
Fasn
Ackr3
Cd34
Tnfrsf12a
TGFbR1_R2
Il4ra
ACVR1_TGFbR
IL6R_IL6ST
IL4R_IL13RA1
Dag1
Pvr
Tnfrsf1a
Il13ra1
Itgb1
Lgals1
Lgals3
ITGAL_ITGB2
Lgals9
Lama5
Mif
Sell
Tnfsf12
Tgfb1
Il4
Il6
Cd226
Tnf
Il13
EBM → iFib
D
upregulated in MPL W515L
ITGAM_ITGB2
Ccr1
Ptprc
C3ar1
ITGA2_ITGB1
ITGAL_ITGB2
Spn
ITGA2B_ITGB3
ITGA6_ITGB1
Cd44
Fcgr2b
Itgal
Itga2
Cxcr4
Ncl
Cd226
Havcr2
Ccr2
Cd69
Itgb1
Cd47
Cxcr2
Cxcl1
Comp
Lgals1
Ccl7
C3
Lgals3
Icam1
Thbs1
Col6a1
Col4a1
Tnxb
Lamb2
Col4a2
Cxcl12
Lgals9
Ptn
Chad
Pvr
Col6a6
iFib → EBM

### Figure S6

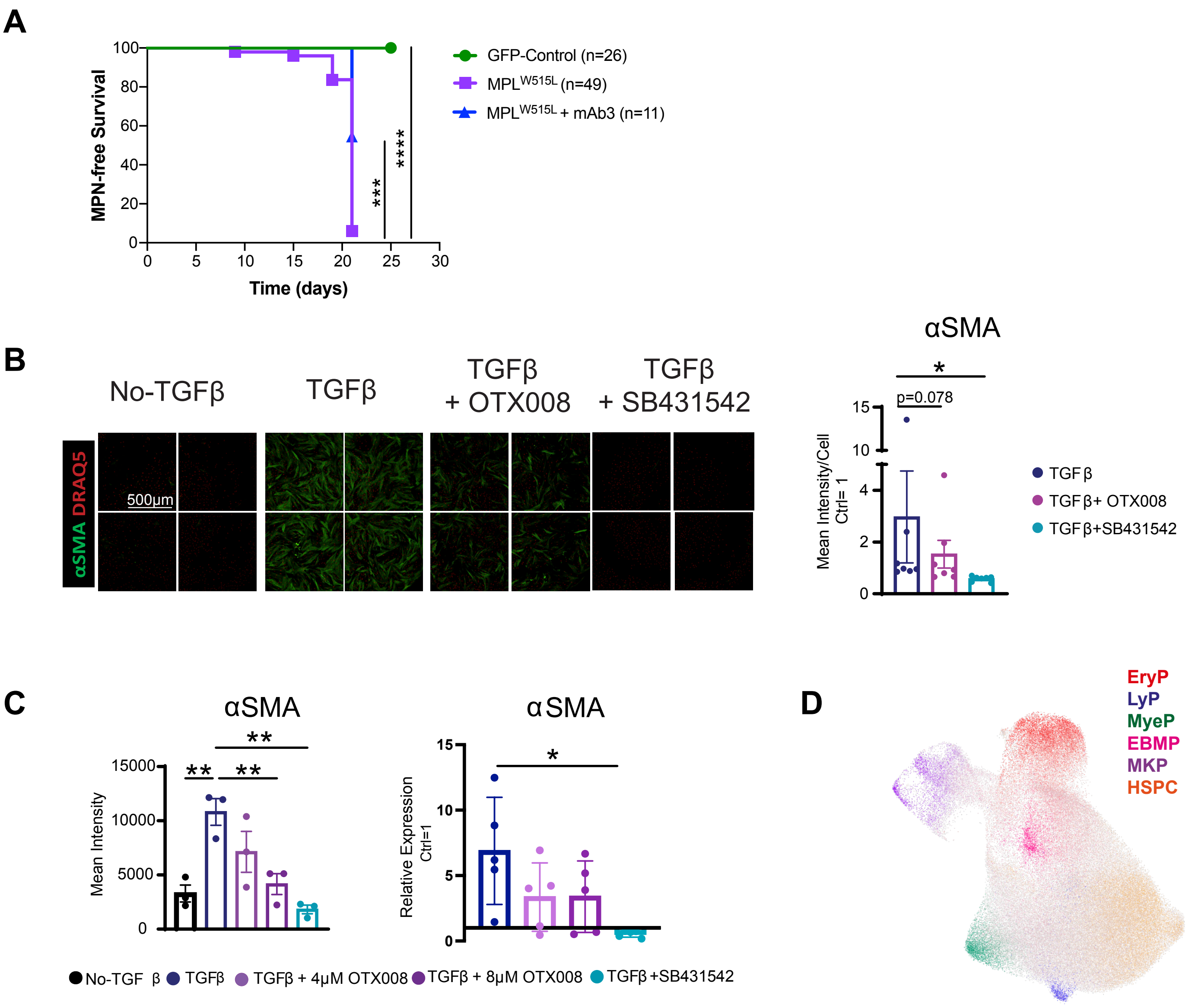

A
MPN-free Survival
Time (days)
GFP-Control (n=26)
MPLW515L (n=49)
MPLW515L + mAb3 (n=11)
***
****
B
No-TGFβ
TGFβ
TGFβ + OTX008
TGFβ + SB431542
αSMA DRAQ5
500μm
αSMA
p=0.078
*
Mean Intensity/Cell Ctrl= 1
TGFβ
TGFβ+ OTX008
TGFβ+SB431542
C
αSMA
Mean Intensity
**
αSMA
Relative Expression Ctrl=1
*
No-TGF β
TGFβ
TGFβ + 4μM OTX008
TGFβ + 8μM OTX008
TGFβ +SB431542
D
EryP
LyP
MyeP
EBMP
MKP
HSPC

#### SUPPLEMENTAL FIGURE LEGENDS

**Supplemental Figure 1. A comprehensive map of myelofibrotic bone marrow, relating to Figure 1.** (**A**) Bar charts showing the body weight of control (n=16) and $MPL^{W515L}$ (n=17) mice, total bone marrow cellularity of 2 femurs and 2 tibias in control (n=20) and $MPL^{W515L}$ mice (n=18), white blood cell count, haemoglobin and platelet counts in control (n=21) and $MPL^{W515L}$ mice (n=17). **p < 0.01, ***p < 0.001, ****p < 0.0001 for unpaired t test with Welch's correction. (**B**) Representative images of haematoxylin & eosin stained spleen from control mice (n=16) and $MPL^{W515L}$ mice. (n=19). Scale bar, 100µm. Red arrows indicate megakaryocytes, and black arrow indicates red pulp. (**C**) Gating strategy for sorting total mononuclear cells, hematopoietic stem/progenitor cells (HSPCs), CD41+ cells and stromal cells for single cell RNA-sequencing. (**D**) Integration of scRNAseq datasets from 3 experiments (Harmony) and removal of doublets (using Scrublet package), prior to louvain clustering. (**E**) Heatmaps showing 5 top differentially expressed genes in each annotated cell type for stromal (left) and hematopoietic cells (right). Abbreviations: BM, bone marrow; MNC, mononuclear cells; Fibro-chondro, fibroblast-chondrocytes; Chondro, chondrocytes; OLC, osteolineage cells; Fibro-osteo, fibroblast-osteoblasts; Fibro, Fibroblasts; MSC, mesenchymal stromal cells; A-endo, arterial endothelial cells; S-endo, sinusoidal endothelial cells; Neutro, neutrophils; GMP, granulocyte-monocyte progenitors; Pro-mono, monocyte progenitors; Mono/MG, monocyte/macrophages; HSC/MPP, hematopoietic stem and multipotent progenitor cells; MK, megakaryocytes; EBM, eosinophil, basophil, mast cells; DC, dendritic cells; B, B cells; T, T cells; Ery, erythrocytes.

**Supplemental Figure 2. Comparison of current dataset to previously published annotations of normal and myelofibrotic bone marrow, and expression of extracellular matrix (ECM) components, relating to Figures 1 and 2.** (**A & B**) Comparison of the stromal (**A**) and hematopoietic (**B**) cells captured to previously published studies by projecting healthy (Baryawano *et al*, blue) and myelofibrotic mouse bone marrow (Leimkuhler *et al*, orange containing stromal cells and Liu *et al,* pink containing haematopoietic cells) onto a reference Uniform Manifold Approximation and Projection (UMAP) plot generated using the cells captured by our study (grey). Abbreviation: Hemat, hematopoietic cells. (**C**) UMAPs showing expression of extracellular matrix factors in stromal cells, broken down into collagens, glycoproteins and proteoglycans. (**D**) As for (C) for haematopoietic cells.

**Supplemental Figure 3. Trans-differentiation of mesenchymal stromal cells and expansion of inflammatory fibroblasts in myelofibrosis, relating to Figure 3.** (**A**) Violin plots showing expression of

*Cxcl12* and *Csf1* in mesenchymal stromal cells (MSC) (left) and fibroblasts (Fibro) (right). Yellow diamond indicates mean value. ***p < 0.001 for Wilcoxon test. (**B**) Uniform Manifold Approximation and Projections (UMAPs) showing expression of canonical MSC marker genes (*Lepr, Adipoq*) and hematopoietic support factors (*Cxcl12 and Kitl*) in MSCs extracted from the main stromal cell dataset. (**C**) Expression of canonical fibroblast marker *Pdgfra and Pdgfrb* on UMAP. (**D**) Pseudotime analysis of fibroblasts using scTour showing that inflammatory fibroblasts (iFib) arise via a separate trajectory from Fib1 cluster (enriched in control mice). The blue cluster indicates the Fib1 cluster.
(**E**) Violin plots showing expression of selected chemokine genes (*Ccl2, Cxcl1 and Kitl*) in fibroblasts in control and $MPL^{W515L}$ mice. Yellow diamond indicates mean value. ***p < 0.001 for Wilcoxon test. (**F**) Violin plot showing expression of *Cxcl5* in iFib vs. all other fibroblasts. Yellow diamond indicates mean value. ***p < 0.001 for Wilcoxon test.

**Supplemental Figure 4. Myelofibrosis megakaryocytes, mast cells and basophils show inflammatory transcriptional programs in myelofibrosis, relating to Figure 4.** (**A**) Uniform manifold approximation and projection (UMAP) plots showing extracted megakaryocytes from the main dataset, confirming high expression of canonical marker genes *Pf4* and *Fli1*. (**B**) Unsupervised clustering of extracted megakaryocytes identified 6 distinct subtypes. (**C**) Identification of megakaryocytes captured from $MPL^{W515L}$ (purple) and control (green) mice. (**D**) Top 5 differentially expressed genes in each megakaryocyte subcluster. (**E**) Heatmap showing gene set enrichment analysis for HALLMARK gene sets in each megakaryocyte subcluster. (**F**) Canonical marker genes for basophils (*Prss34*), mast cells (*Tpsb2*) and eosinophils (*Prg2)* respectively shown on EBM cell UMAP.

**Supplemental Figure 5. Up-regulated receptor-ligand interactions in myelofibrotic bone marrow, relating to Figure 5.** (**A** & **B**) Interaction network of **(A)** TNF and (**B**) IL4 signalling pathway indicating EBM cluster is the key ligand resource (sender) for both pathways. (**C** & **D**) Circus plot depicting upregulated interaction pairs in $MPL^{W515L}$ vs control mice between EBM cluster and iFib cluster highlighting the enrichment of *Lgals1* interactions.

**Supplemental Figure 6. Inhibition of galectin 1 signalling reduces myelofibrosis phenotype *in vitro* and *in vivo,* relating to Figure 7. (A)** Kaplan–Meier curve showing MPN-free survival (defined by blood parameters and spleen size) for control mice (n=26), $MPL^{W515L}$ mice (n=49) and anti-Gal1 mAb3 treated $MPL^{W515L}$ mice (n=11). *** p < 0.001, **** p < 0.0001 for Gehan-Breslow-Wilcoxon test. **(B)** TGFβ-induced fibroblast to myofibroblast transition assay using human bone marrow stromal cells treated with TGFβ alone ± the galectin 1 inhibitor OTX008 and the TGFβ inhibitor SB431542. Representative images shown (left) from high-throughput 384-well plate assay. Chart (right) shows Mean Fluorescence Intensity per cell for αSMA normalized to the no-TGFβ control ± SEM (n=7). *p < 0.05 for wilcoxon matched pairs signed rank test. (**C**) αSMA protein quantification by immunofluorescence staining intensity and RT-PCR for gene expression of human bone marrow organoids treated with TGFβ to induce organoid fibrosis ± OTX008 or SB431542. n=5-8 organoids from 3 independent experiments. *p < 0.05, ** p < 0.01 for one-way ANOVA. (**D**) Identification of eosinophil, basophil and mast cell progenitors (EBMP) within a published dataset of >120,000 CD34+ Lineage negative hematopoietic stem/progenitor cells from patients with myelofibrosis and age-matched healthy donors.
